# Structural Basis for Stereoselective Dehydration and Hydrogen-Bonding Catalysis by the SAM-Dependent Pericyclase LepI

**DOI:** 10.1101/491761

**Authors:** Yujuan Cai, Yang Hai, Masao Ohashi, Cooper S. Jamieson, Marc Garcia-Borras, K. N. Houk, Jiahai Zhou, Yi Tang

## Abstract

LepI is an *S*-adenosylmethionine (SAM)-dependent pericyclase that catalyzes the formation of 2-pyridone natural product leporin C. Biochemical characterization showed LepI can catalyze the stereoselective dehydration to yield a reactive (*E*)-quinone methide which can undergo a bifurcating intramolecular Diels-Alder (IMDA) and hetero-Diels-Alder (HDA) cyclization from an ambimodal transition state, and a [3,3]-retro-Claisen rearrangement to recycle the IMDA product into leporin C. Here we solved the X-ray crystal structures of SAM-bound LepI, and in complex with a substrate analog, the product leporin C, and a retro-Claisen reaction transition-state analog to understand the structural basis for the multitude of reactions. Structural and mutational analysis revealed how Nature evolves a classic methyltransferase active site into one that can serve as a dehydratase and a multifunctional pericyclase. Catalysis of both sets of reactions employ His133 and Arg295, two active site residues that are not found in canonical methyltransferases. An alternative role of SAM, which is not found to be in direct contact of the substrate, is also proposed.

## INTRODUCTION

Pericyclic reactions are among the most powerful synthetic reactions to make multiple regioselective and stereoselective carbon-carbon bonds and carbon-heteroatom bonds.^1^ Despite their prevalence in organic synthesis, only a handful of naturally occurring enzymes have been characterized to catalyze pericyclic reactions and related [4+2] cycloadditions.^2^ Structural characterizations of a few of these enzymes, including chorismate mutase (CM), isochorismate lyase, precorrin-8x methyl mutase, SpnF, etc., showed Nature has evolved a variety of protein folds divergently to accelerate the pericyclic reactions, and control the regio- and stereoselectivity.^3–11^

Recently we discovered a multifunctional *S*-adenosyl-L-methionine (SAM) dependent *O*-methyltransferase (OMT)-like pericyclase, LepI, that catalyzes a cascade of reactions starting from the 2-pyridone alcohol **2** to form the dihydropyran-containing fungal natural product leporin C (**10**) (**Figure 1**).^12^ In the absence of LepI, the alcohol **2,** which is derived from the ketoreduction of **1**, can dehydrate into either the *(E)-* or (*Z*)-quinone methide **3** or **4,** respectively. The reactive **3** and **4** can undergo intramolecular Diels-Alder (IMDA) cycloaddition to yield the *endo* products **9** and **6**, respectively, as well the *exo* adducts **8** and **5,** respectively. The inverse-electron demand hetero-Diels Alder (HDA) cycloaddition of **3** and **4**, on the other hand, affords the desired product **10** and the diastereomeric **7**, respectively. Quantum mechanics (QM) calculations revealed that the activation energies for the formation of these six products from either **3** or **4** are comparable, consistent with **10** being only a minor product in the absence of LepI. We showed that in the presence of LepI, **2** is completely converted to **10**.^12^ We also demonstrated computationally that the *endo* IMDA and HDA reactions starting from **3** goes through an ambimodal transition state (**TS-1**) and that a post-transition-state bifurcation leads preferentially to the IMDA product in the absence of enzyme catalysis. Therefore, LepI must catalyze the diastereoselective-dehydration of **2** to form only the (*E*)-quinone methide **3**; followed by facilitating the *endo* transition state (**TS-1**) from **3,** while suppressing the *exo* transition state (**TS-3**). The active site of LepI can also alter the dynamics of the bifurcating potential energy surface from **TS-1** to favor the HDA product **10** as suggested by computation.^12^ Furthermore, we demonstrated LepI can catalyze the retro-Claisen reaction of converting **9** into **10** via the [3,3]-sigmatropic retro-Claisen rearrangement **TS-2**, as a means of kinetically recycling the IMDA shunt **9** to product **10**. While our biochemical and computational studies unveiled these unexpected roles of an enzyme that has sequence homology to an OMT, the mechanistic details of how catalysis and selectivity are achieved by LepI were not known.

**Figure 1.**
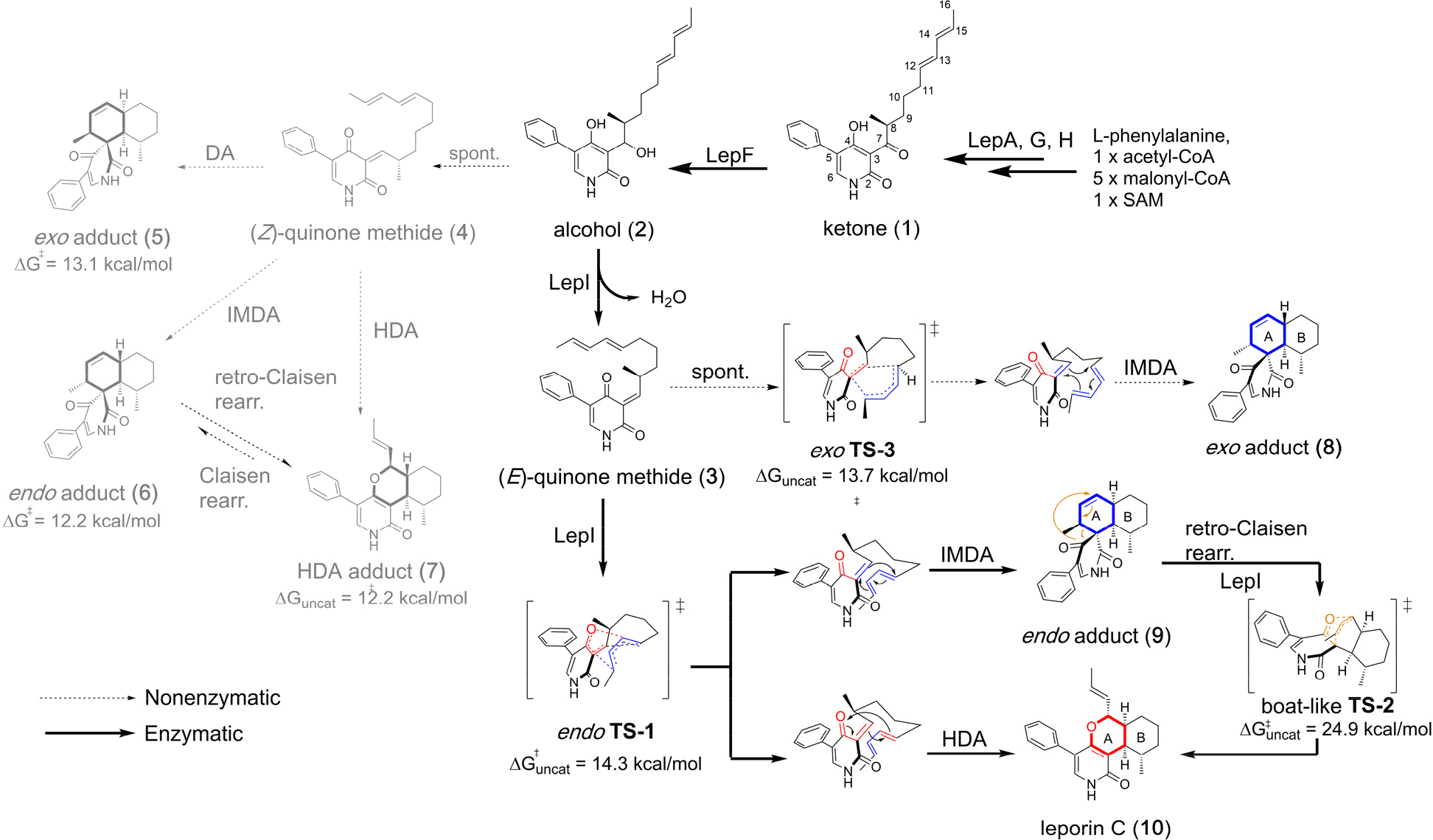
Leporin C biosynthesis pathway highlighting LepI catalyzed reaction cascade. In the absence of LepI, spontaneous dehydration of alcohol **2** yields a (*E/Z*)-mixture of quinone methide (**3**/**4**), which nonenzymatically form Diels-Alder and hetero Diels-Alder adducts.

Here we report a collection of X-ray crystal structures, which, combined with computational studies, uncover the origins of catalysis and stereoselectivity of LepI. We obtained X-ray crystal structures of *holo*-LepI i) bound with SAM; ii) complexed with the dehydration reaction substrate analogue **1**; iii) complexed with product leporin C **10**; and iv) complexed with the retro-Claisen reaction substrate analogue **8**. Our structures provide insights regarding the molecular mechanism of LepI catalysis, including the *E*-stereoselectivities of dehydration, the *endo*-stereoselectivity of the IMDA/HDA reactions, and the rate enhancement of the retro-Claisen reaction. A possible divergent evolutionary relationship with SAM-dependent OMTs and a new role of SAM in catalyzing pericyclic reactions are also proposed.

## RESULTS

### Structure of LepI

The structure of LepI was determined by single anomalous diffraction on a Se-Met derivative crystal diffracting to 2.58 Å resolution, and the phase was extended to a native crystal with 2.14 Å resolution by molecular replacement (**Supplementary Table 1**). The overall tertiary structure of LepI comprises an *N*-terminal dimerization domain (residues 1-154) and a *C*-terminal catalytic domain (residues 155-387) which adopts the typical α/β Rossmann fold observed in all class I SAM-dependent methyltransferase (**Figure 2a**).^13^ In the asymmetric unit, two LepI monomers intertwine through the *N*-terminal dimerization domains with a dimer interface of ~5,000 Å^2^ surface area per monomer as calculated by PISA (**Figure 2b**).^14^ Such intimate homodimer architecture is similar to that of many OMTs.^2^ Unique to LepI dimer is a domain-swapped four-helix bundle at the *N*-terminus. This leucine-rich coiled-coil enables LepI dimers to pack against each other and form a dimer of dimers, burying ~1,800 Å^2^ surface area between two dimers (**Supplementary Figure 1**). In comparison, the oxaline *O*-methyltransferase OxaC,^15^ the closest homologue to LepI as identified by Dali,^16^ also harbors a similar four-helix bundle at the *N*-terminus but it is not domain-swapped and does not further dimerize. The tetramer oligomerization of LepI is consistent with sedimentation velocity experiment (**Supplementary Figure 2**), which showed LepI exists in an equilibrium between dimer and tetramer in solution.

**Figure 2.**
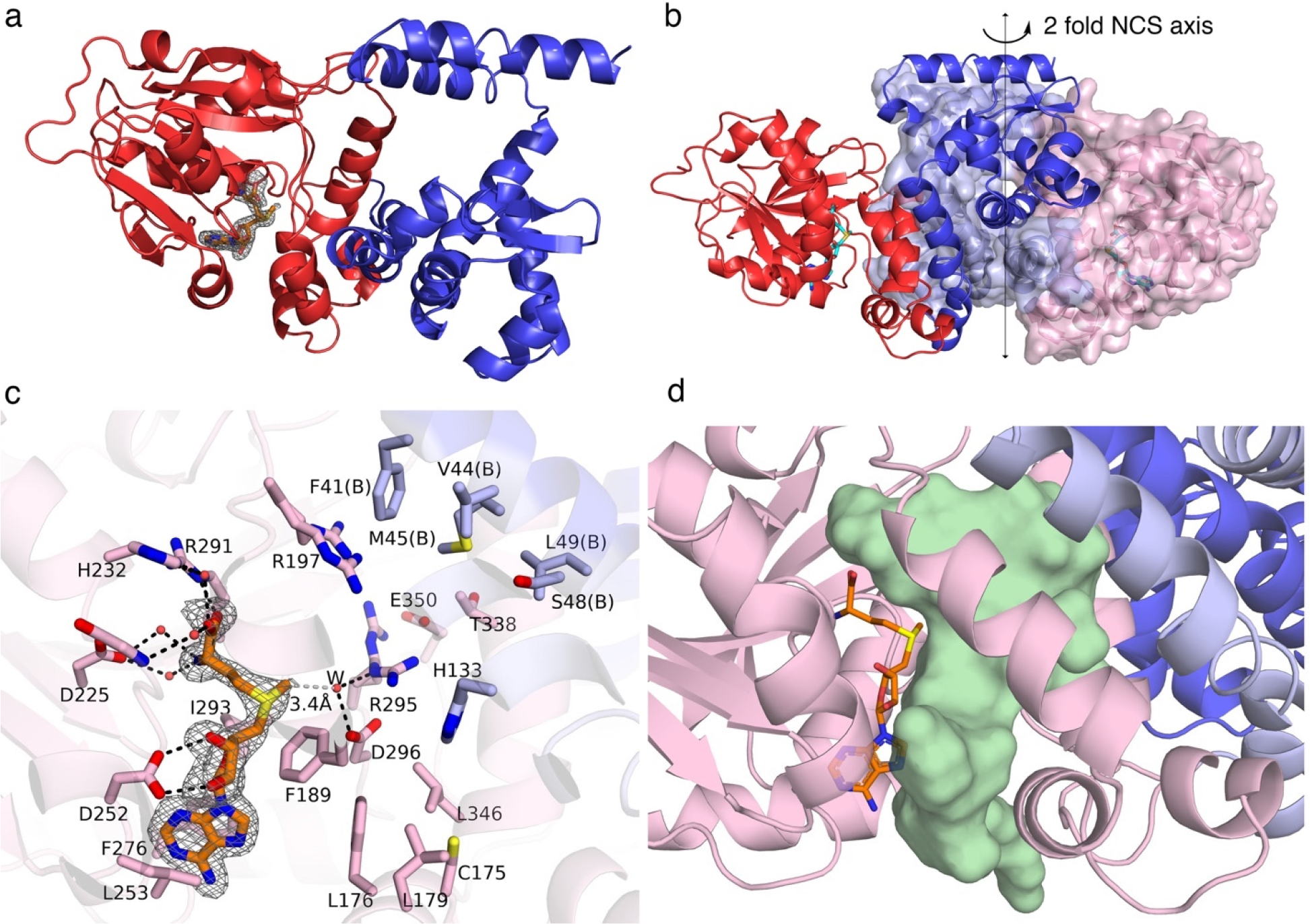
LepI structure and the SAM binding site. (a) The overall tertiary structure of LepI is shown in cartoon model, with the *N*-terminal dimerization domain and *C*-terminal catalytic domain colored in blue and red, respectively. Simulated-annealing omit map (grey mesh, contoured at 2.5 σ) indicates binding of SAM at the canonical SAM binding site. (b) Intimate LepI homodimer featured by an intertwined dimer interface. (c) Close-up view of SAM binding site. Hydrogen bond interactions are indicated with black dashes. (d) Next to SAM is a large substrate binding cavity and its entrance tunnel (shown together as green surface). The volume of cavity was calculated by using POCASA.^19^

The cofactor SAM is copurified with LepI with >90% occupancy when it is overexpressed in *E. coli*.^12^ No exogenous SAM was supplemented during protein purification and crystallization steps. Electron density unambiguously delineates the binding of SAM in each LepI monomer at the canonical SAM binding site (**Figure 2c**). LepI makes numerous contacts with SAM that are observed in the bona fide OMTs: the α-amino and α-carboxylate groups are recognized through direct and water-mediated hydrogen bond networks; the diol group from ribose ring donates two hydrogen bonds to D252; the adenine ring is sandwiched between L253 and F276 involving van der Waals and π…π interactions. Notably, the sulfonium methyl group is making an unconventional CH…O hydrogen bond with D296, mediated by a water molecule (designated as **W**, the distance between SAM sulfonium methyl carbon and **W** is 3.4 Å). Such CH…O hydrogen bonds involving SAM sulfonium methyl group are prevalent in SAM-dependent methyltransferases and are believed to be important for SAM binding specificity and facilitating methyl transfer reaction.^17^ However, water molecules have rarely been observed as immediate hydrogen bond acceptor with SAM.^18^ The water molecule **W** is also making a hydrogen bond with R295, which resides at the center of a ~500 Å^3^ large cavity. This putative
active site cavity is outlined by both the catalytic domain (monomer A) and the dimerization domain from the other monomer (monomer B) within the homodimer unit **(Figure 2d)**. Adjacent to the adenine ring of SAM is a wide substrate entrance tunnel, whereas on the backside of the cavity at the domain interface, multiple water-filled tunnels were also found to connect the cavity to bulk solvent (**Supplementary Figure 3)**. Such organization implies that the dimerization event is obligatory for LepI function and each homodimer is an integral and minimal functional unit.

### Structure of LepI-substrate analogue ketone (1) complex (LepI-SAM-1)

To understand how LepI recognizes and orients the substrate alcohol **2** and achieves (*E*)-selective dehydration to yield **3**, we cocrystallized LepI with ketone **1**, which serves as an unreactive substrate analogue. This pseudo enzyme-substrate (Michaelis) complex was determined to 1.70 Å resolution. Electron density clearly reveals the binding of **1** in the proposed substrate binding site adjacent to SAM, as well as an ethylene glycol molecule serendipitously sitting on top of the pyridone ring (**Figure 3a**). The atomic resolution of the electron density also allows us to determine the absolute stereochemistry of the chiral C8 of **3** (*S* configuration).

**Figure 3.**
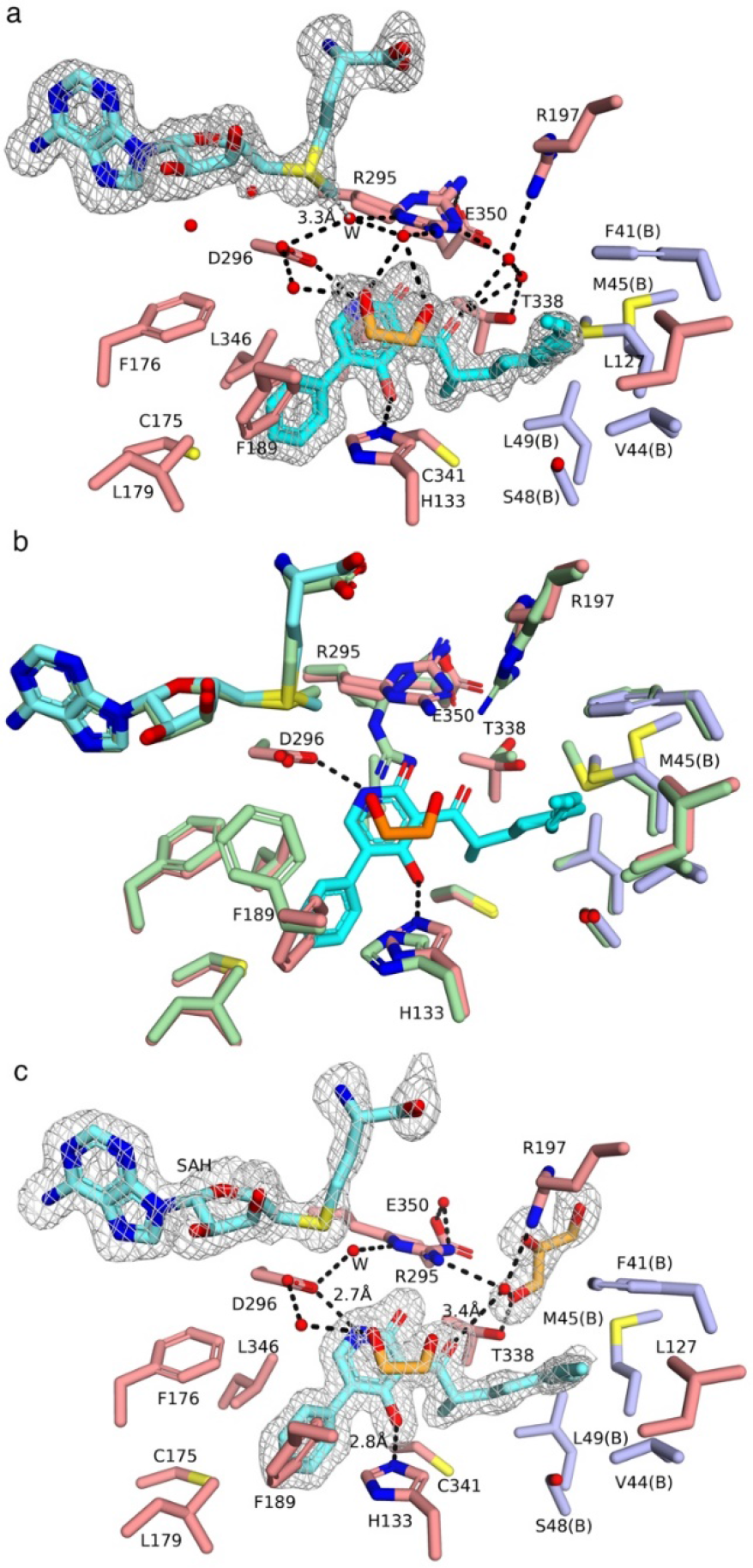
Crystal structure of LepI pseudo enzyme-substrate complex. Simulated-annealing omit maps are shown in black mesh and contoured at 3.0 σ Hydrogen bond interactions are indicated by black dashed lines. (a) Crystal structure of LepI-SAM- **1** (pseudo enzyme-substrate complex). Note that two rotamers of R295 and M45(B) are observed, and two conformations of diene (*s*-trans and *s*-cis) are modeled according to the electron density. Residues from monomer A are colored in salmon, whereas residue from monomer B are colored in light blue and labeled with B in parenthesis. (b) Superposition of LepI-SAM-**1** complex with unliganded LepI-SAM (colored in pale green). Substantial conformational changes are observed with F189, R295, R197, and M45(B). (c) Crystal structure of LepI-SAH-**1** ternary complex with a glycerol in the water exit tunnel.

The overall LepI-SAM-**1** structure is very similar to the ligand-free structure with an r.m.s.d. of ~ 0.18 Å for 770 C_α_ atoms. However, local structural changes were triggered upon substrate binding (**Figure 3b)**. The 5-phenyl ring of **1** is locked into a small hydrophobic pocket, and confers favorable π…π interaction with H133 and F189, which caused rotation of F189 side chain. The pyridone ring is recognized via hydrogen bond interactions with H133 and D296 through the 4-OH group and amide carbonyl, respectively.

Unexpectedly, instead of being frozen in the near-attack conformation (NAC) for the cycloaddition step immediately following dehydration, the diene alkyl chain is captured in an extended conformation projecting into a hydrophobic cleft mainly surrounded by residues from monomer B within the homodimer (**Supplementary Figure 4)**, and accommodation of this linear alkyl chain caused a ~3 Å shift of M45(B) side chain. Meanwhile, the space left for the diene chain at the NAC is now occupied by the ethylene glycol molecule. The electron density reveals that the diene moiety adopts alternate conformation (*s*-trans and *s*-cis). In both conformations the diene could confer favorable π…π interaction with F41(B).

The C7 carbonyl group is oriented in parallel with the amide carbonyl facing towards the aforementioned solvent-filled channel, and forms an extensive water hydrogen bond network mediated by residues T338, R197, R295, and E350. Since the C7 carbonyl group closely mimics the leaving alcohol group of substrate **2** in the transition state of the elimination reaction, such orientation dictates that substrate **2** must undergo *anti*-elimination exclusively yielding (*E*)-quinone methide **3**, which explains our previous biochemical characterization. The water hydrogen bond network helps to protonate the leaving alcohol group. The byproduct water may be transiently trapped by the nearby polar residues, and then be eventually exported to bulk solvent through the solvent releasing channel.

Even though the substrate binding pocket is in proximity to cofactor SAM, no direct contact was observed between SAM and ketone **1** (the distance between SAM sulfonium methyl carbon and the ketone **1** amide carbonyl oxygen atom is about 6.8 Å). Instead, the water molecule **W** remains to bridge the SAM sulfonium methyl group with D296 and R295. A structure of LepI-SAH-**1** ternary complex (1.84 Å resolution) was also determined during the course of the study, in which *S*-adenosyl-L-methionine (SAH), instead of SAM, was identified in the active site (**Supplementary Figure 5**). Even though SAH lacks the sulfonium methyl and formal positive charge compared to SAM, the overall structure is essentially identical to the LepI-SAM-**1** complex, including the active site organization, waters, and substrate binding mode, except that the diene exclusively adopts the *s*-trans conformation; and an extra glycerol molecule was found at the exit of the solvent-releasing channel. The similarity between these two complexes argues that any difference reflected on catalysis exerted by SAM *vs* SAH as observed before, must not result from a structural role or allosteric effect; and should be attributed to the hydrogen bonding and positively charged nature of SAM *vs* SAH.

### Structure of LepI enzyme-product (10) complex (LepI-SAM-10)

To gain more insight of how LepI catalyzes cycloaddition, we next determined the crystal structure of LepI-SAM complexed with leporin C (**10**) to 1.78 Å resolution. As expected, no major overall conformational changes of LepI were induced upon binding of the product. The crystal structure of enzyme-product complex reveals that **10** is accommodated in a similar manner as seen in the LepI-SAM-**1** complex (**Figure 4a**) through hydrophobic effect and hydrogen bond interactions. Notably, residue R295 now adopts a different conformation to form a bifurcated hydrogen bond with the amide carbonyl of **10**. The amide carbonyl concurrently accepts a hydrogen bond donated from a water molecule (**W’**), which likely originates from the leaving alcohol group after dehydration. This water molecule also participates in forming the hydrogen bond network at the solvent releasing channel. The direct hydrogen-bond interaction between the amide carbonyl with guanidinium group of R295 is reminiscent of the electrostatic catalysis predicted previously by our computational study: a sulfonium ion or ammonium ion interacts with the amide carbonyl in the ambimodal **TS-1**.^12^ Such electrostatic catalysis was predicted to decrease the C-O bond length and increase the C-C bond length in **TS-1**. This contributes to the observed preference for the HDA reaction versus *endo*-mode IMDA in the presence of LepI as compared to non-enzymatic reaction in water. In addition to the amide carbonyl, a lone pair from cyclic ether oxygen is shown to accept a hydrogen bond from the side chain of H133. Considering that the general base H133 is inevitably protonated after dehydration in the reaction cascade, H133 is most likely present as the imidazolium form, which makes it as an ideal hydrogen bond catalyst for the following pericyclic reactions. The positively charged imidazolium ring of H133 maintains hydrogen bonding with the C4-carbonyl of the quinone methide **3** and **TS-1** to enhance electrophilicity of the pyridone ring. In the [3,3]-sigmatropic retro-Claisen rearrangement, this interaction also stabilizes the developing partial oxyanion in the **TS-2**. Together, hydrogen bonds from R295 and H133 to C2 and C4 carbonyl groups activate the quinone methide **3** and the retro-Claisen substrate **9** by stabilizing the polarized transition state.

**Figure 4.**
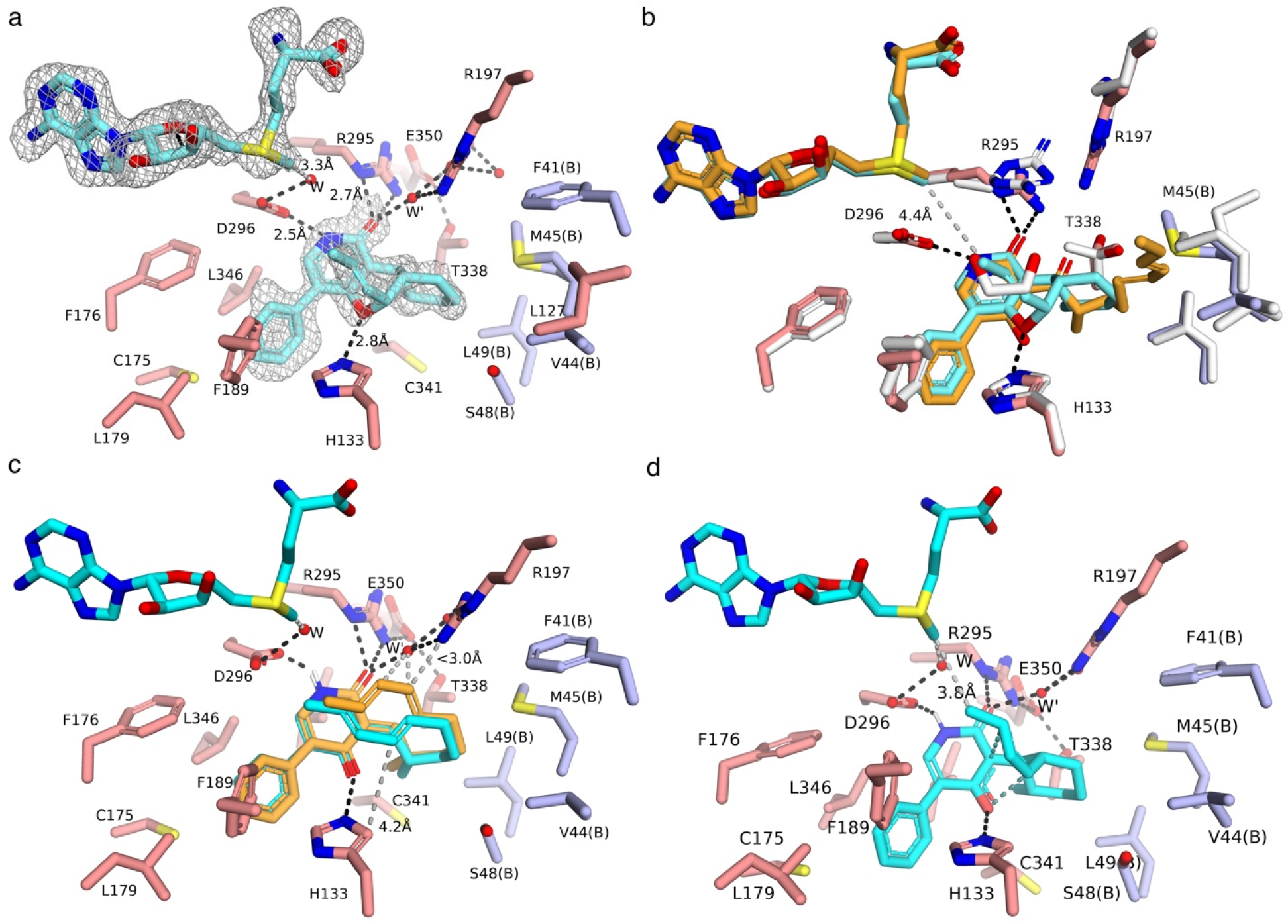
Crystal structure of LepI enzyme-product (10) complex. Simulated-annealing omit map is shown in black mesh and contoured at 3.0 σ SAM and ligands are colored in cyan. Hydrogen bond interactions are indicated by black dashed lines. (a) Crystal structure of LepI-SAM-**10**. (b) Superposition of LepI-SAM-**10** with LepI-SAM-**1** (residues colored in white, SAM and ligand are colored in orange). Noticeable conformational changes are R295, R197, T338, and M45(B). (c) TS-1 (cyan) is docked into LepI. The distance (4.2 Å) between H133 N_ε_ and TS-1 C14 is indicated by grey dashes. In comparison, the competing TS-3 (orange) in nonenzymatic reaction is modeled into LepI. (d) TS-2 (cyan) is docked into LepI. The breaking C-C bond and forming C-O bond during the transition state are indicated with dashed lines (cyan).

Superimposing the LepI-SAM-**10** complex onto LepI-SAM-**1** complex reveals that cycloaddition of the diene chain causes the conformational change of M45(B), which swings back to the original position as seen in the unliganded structure (**Figure 4b**). Note that the previously observed ethylene glycol molecule overlays well with C14-C16 of **10**, which illuminates how the linear portion of quinone methide **3** rearranges to the NAC preceding **TS-1**: the diene alkyl chain of quinone methide **3** is induced by the enzyme binding site to rotate and fold on top of the 2-pyridone ring, leaving the C8-C13 region preorganized into a chair-like conformation with C8 methyl group in the equatorial position and the diene moiety in *s*-cis conformation. Accordingly, we docked the ambimodal *endo* **TS-1**, and the [3,3]-sigmatropic retro-Claisen rearrangement **TS-2** into the LepI-SAM-**10** structure omitting ligand **10** (**Figure 4c,d**). Including the two well-defined water molecules (**W** and **W’**) and cofactor SAM in the structure greatly facilitated docking the ligands in the productive poses. This supports the hypothesis that they may play important structural roles in defining the active site binding environment.

The basis for LepI diastereoselectivity (*endo* vs *exo*) in the IMDA/HDA reaction was gained by modelling the hypothetical LepI-SAM-**TS-3** complex with the assumption that the 5-phenyl group and pyridone ring are bound as in **TS-1** to maintain similar interactions with LepI (**Figure 4c**). In order for the *exo* **TS-3** to bind, the hydrophobic diene group (C13-C14) is positioned on the hydrophilic side of the active site, and potentially causes steric clashes with water **W’** and R295; whereas in the *endo* **TS-1** model, the diene is positioned on the hydrophobic side of the binding site. Thus, the diastereoselectivity likely arises from preference of the diene to avoid the hydrophilic “wall” constituting of polar residues (D296, R295, R197, T338) and **W’**.

### Structure of LepI enzyme-intermediate analogue (8) complex (LepI-SAM-8)

The proposed catalyzed transition structures are further corroborated by the LepI-SAM-**8** ternary complex structure (1.66 Å resolution), where the unreactive spirocyclic nonenzymatic *exo* **TS-3** product **8** served as a surrogate to mimic the diastereoisomer spirocyclic **9** when bound to the enzyme (For structural characterization of **8**, see **Supplementary Table 3**). Key hydrogen bond interactions are retained in recognizing **8**, except the water **W’** (**Figure 5**). The disappearance of **W’** in LepI-SAM-**8** complex is due to apparent steric clash between **W’** and C14 of **8**, which is derived from the *exo* **TS-3**. This observation further supports the conclusion based on docking experiments that the presence of **W’** disfavors formation of **TS-3**.

**Figure 5.**
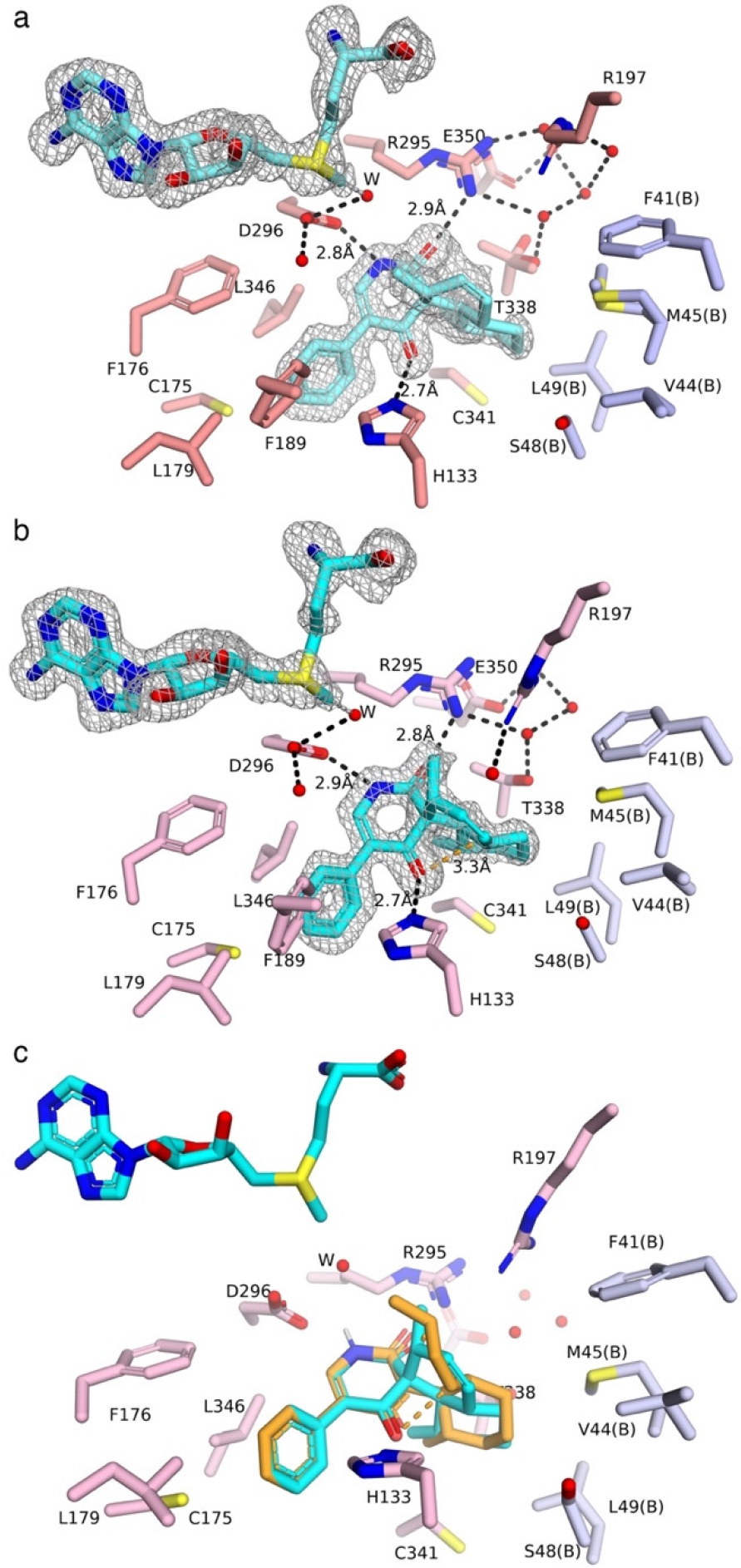
Crystal structure of LepI enzyme-intermediate analogue (8) complex. Simulated-annealing omit map shown in black mesh and contoured at 3.0 σ. Hydrogen bond interactions are indicated by black dashed lines. (a) Conformer A of ligand **8**. (b) Conformer B of ligand **8.** The shortened distance between carbonyl oxygen atom and C13 in (b) is indicated by an orange dashed line. (c) Superposition of conformer B of ligand **8** with TS-2 shows remarkable similarity.

Two alternative conformations were observed for the spiro-decalin ring of **8**. In both conformations, the B ring adopts the chair conformation. However, in monomer A (**Figure 5a**), the A ring adopts a “relaxed” twisted-boat conformation resembling that in the crystal structure of **8** determined by single crystal X-ray analysis (**Supplementary Figure 6**); whereas in monomer B (**Figure 5b)**, the A ring is found in a thermodynamically less stable (~1 kcal/mol according to our DFT calculations) half-chair conformation bringing the C=C double bond closer to the C4 carbonyl group, resulting in a distance of 3.3 Å between the carbonyl oxygen and C13 atom from the double bond. This less stable half-chair conformation captured *in crystallo* resembles that of the retro-Claisen reaction transition state **TS-2** instead of the substrate **9** when docked in LepI (**Figure 5c**). In general, an enzyme favors the binding of a transition state-like structure over a substrate-like structure, and the intermediate analogue **8** serves as a transition state (**TS-2**) analogue when bound to LepI.^20^ LepI does not promote a [3,3]-sigmatropic retro-Claisen rearrangement of **8** because the requisite transition state TS-4 has an intrinsically high activation energy barrier (10.7 kcal/mol higher compared to that from **9** to **TS-2**). However, the binding of the half-chair conformer of **8** by LepI implies that the LepI active site is well-tailored for stabilizing the retro-Claisen rearrangement transition state **TS-2** (**Supplementary Figure 6c**).

### Catalytic mechanism of LepI

To verify our interpretations of likely modes of catalysis, we performed site-directed mutagenesis on the implicated residues and examined the enzyme activity in *A. nidulans*. In the presence of upstream enzymes that produce the alcohol **2**, a fully active LepI yields only the product **10**, while a completely inactive LepI (or absence of LepI) leads to a mix of **5-10** (**Figure 6a**). Consistent with our structural conclusion, removal of the essential general base H133 is detrimental to LepI activity. All four mutations (H133A, H133F, H133Q, H133N) completely abolished enzyme activity (**Figure 6a**). As observed in the LepI-SAM-**1** structure, R295 may act as a general acid and indirectly protonate the leaving alcohol. Substitution of R295 with Ala/Gln/Phe/Tyr significantly compromised the dehydration activity, but activity can be effectively rescued by R295H/K/N. Mutation of other residues structurally implicated in the protonation step during dehydration (T338A, T338S, R197A, and R197K) did not affect the activity, which suggest they are not essential in catalysis. Residue D296 plays a major role in substrate binding and polarizing the substrate in order to lock the substrate in the 2-pyridone form versus its tautomeric 2-hydroxypyridine form. Substitution of residue D296 with Ala led to notable increase in shunt products, and the activity can be restored by D296N and D296E, both of which can accept similar hydrogen bonds from pyridone NH. Even though substantial conformational change was observed on M45 accompanying intermediate cycloaddition, M45A mutation did not affect the yield *in vivo* at all, which strongly suggests that folding of the diene chain of quinone methide **3** to the NAC preceding **TS-1** is spontaneous.

**Figure 6.**
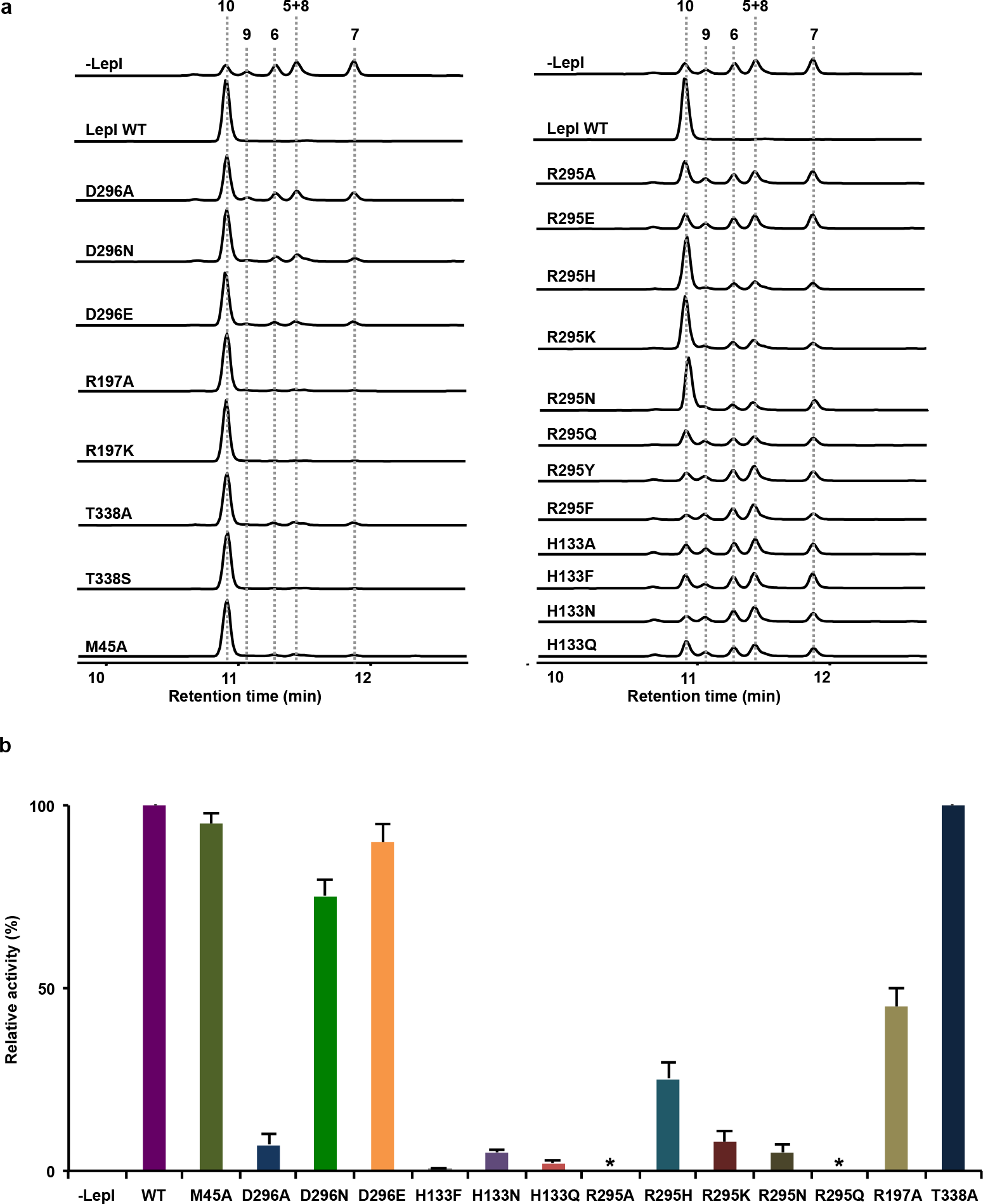
LepI Mutant activity. (a) *In vivo* activity of LepI or mutants to synthesize leporin C (**10**) starting from the alcohol substrate **2. 2** is synthesized by upstream biosynthetic enzymes, for details of pathway see Ref 12. (b) *In vitro* retro-Claisen rearrangement activity using **9** as the substrate. Asterisks indicate mutants with no measurable activity.

To examine the contribution from these active site residues to the catalysis of retro-Claisen rearrangement reaction, we assayed the enzymatic activity of the purified mutants using **9** as the substrate in vitro. Consistent with in vivo result, mutation of non-catalytic residues (T338A, M45A) did not affect the pericyclic activity. Alanine substitution for the substrate binding residue D296 causes a 10-fold loss of activity, but the activity can be restored to 75% and 90% of wild-type by D296N and D296E mutants, respectively. In contrast, mutation of the hydrogen bonding catalyst H133 significantly compromised the pericyclic activity: H133F decreased the activity by 200-fold; moreover, replacing H133 by other neutral hydrogen bond donors (e.g. H133Q and H133N) did not restore the activity, which demonstrates the necessity of the positively charged imidazolium group for this pericyclic reaction. Ala and Gln substitution at R295 completely abolished the activity. Substitution by Asn and Lys at R295 retained 10% activity of wild-type, whereas His substitution restores the activity to 30% of wild-type. The successful partial rescue by the R295H mutant suggests that the imidazolium ring can be stabilized by the nearby second-shell residue E350 and donates a favorable hydrogen bond to the amide carbonyl to catalyze the retro-Claisen rearrangement. R197, which makes a water mediated (W’) hydrogen bond to the same amide carbonyl, plays a minor catalytic role in this reaction, as R197A mutant retained 50% activity.

Taken together, this structure-activity relationship study provides mechanistic insights into LepI catalysis **(Figure 7)**. The dehydration step likely proceeds via the E1-cb mechanism: H133 acts as the key general base to deprotonate the 4-OH group and stabilize the corresponding enolate anion intermediate, while the leaving hydroxyl group is ejected *anti*-periplanar and protonated by water molecules from the water hydrogen bond network, a process that is facilitated by R295. The leaving water molecule (**W’**) could be transiently trapped by these residues and maintain a hydrogen bond with the amide carbonyl. The lack of a dedicated general acid residue to facilitate dehydration presumably is to prevent the newly formed highly reactive quinone methide intermediate *(E)-***3** from being quenched by **W’**, since a good general acid residue could act in principle as a general base in the reverse reaction. The residue D296 is important for substrate binding and favoring the 2-pyridone tautomer. Overall, the dehydration specificity (1,4-*anti*-elimination) was accomplished through positioning the general base and acid *in trans* configuration, and trapping the linear alcohol substrate **2** in the corresponding *trans*-conformer.

**Figure 7.**
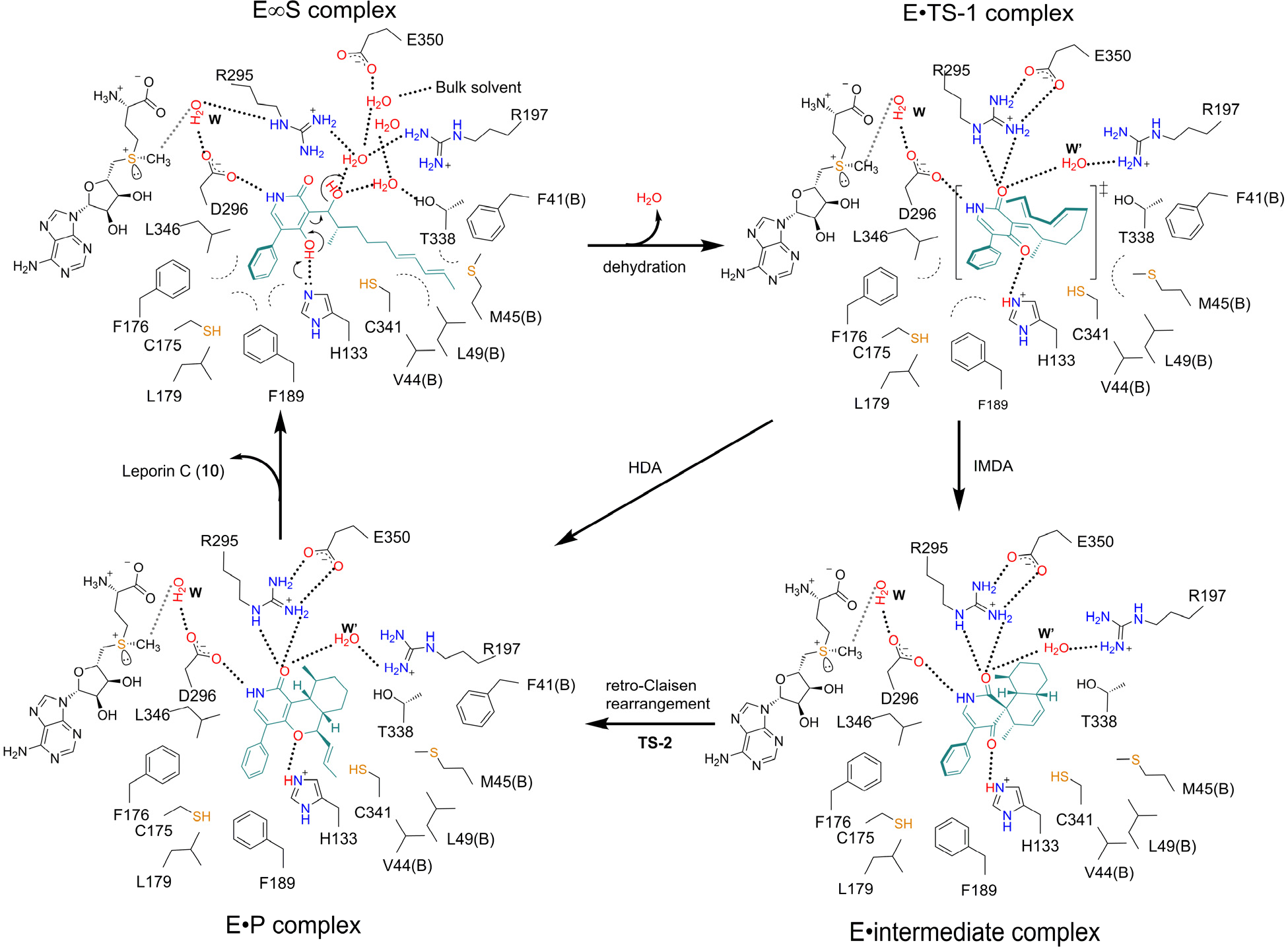
Proposed catalytic mechanism of LepI. Dashed lines indicate hydrogen bonds.

For the following cycloaddition reaction, spontaneous bond rotation of diene is driven by filling up the void volume over pyridone ring, and better shape-complementarity in the active site when poised for reaction. The diastereoselectivity was achieved by aligning hydrophilic residues at the *exo* side which in turn favors the *endo* **TS-1** (hydrophobic effect). Residues H133 and R295 act as hydrogen bond catalysts to lower the transition state energy barrier and stabilize both **TS-1** and **TS-2**. In particular, the highly polarized transition state **TS-2** (i.e. the oxyanionic 4-carbonyl oxygen) can be electrostatically stabilized by the positively charged imidazolium side chain of H133. The cofactor SAM is required in defining the active site shape, but not directly interacting with the substrate. The catalytic role of SAM during the reaction as shown by previous biochemical experiments, and the necessity of a charged analog to effect catalysis is indicative of the electrostatic role of the SAM sulfonium moiety for catalysis, but it is too remote for direct hydrogen bonding.

## DISCUSSION

A common theme found in pericyclases^2^ that catalyze [4+2]-additions is that cyclization follows an initial priming/deprotection reaction which activates the substrate to create/unmask the reactive functional groups.^21^ This priming/deprotection reaction may be catalyzed by a separate enzyme (e.g., tandem reaction catalyzed by SpnM and SpnF); or the cyclase itself (e.g. solanapyrone synthase), which generates the reactive intermediate *in situ*. The second scenario is most often encountered when the liberated intermediate is too reactive and readily forms undesired cyclized products without enzymatic control. Therefore, by devising such a cascade scheme, the unstable intermediate generated *in situ* will undergo cyclization immediately inside the cyclase.

LepI also follows this principle, and our studies here provided the structural basis and mechanistic insight for LepI catalysis, such as 1) positioning the general base (H133) and general acid (water facilitated by R295) in the *trans*-configuration to restrict the substrate **2** to undergo *anti*-elimination yielding (*E*)-quinone methide **3**; 2) setting the “amphiphilic” active site to favor the *endo*-conformation during cycloaddition; 3) repurposing the two cationic residues (H133 and R295) as hydrogen bonding catalysts to lower the transition state barrier and to electrostatically stabilize the polarized transition state; 4) repurposing the SAM normally functional in methyl transfers to in a likely electrostatic role for catalysis.

In both natural and artificial enzymes that catalyze Diels-Alder reactions, it has been found that a strategy for rate enhancement is to donate specific hydrogen bonds to the dienophile in order to lower its LUMO energy^22,23^ and to stabilize the polarized transition state. In LepI catalysis, H133 and R295 are vivid examples as hydrogen bond catalyst to donate hydrogen bonds to both the amide carbonyl and quinone methide carbonyl simultaneously. Such hydrogen bonding effectively lowers the LUMO energy of quinone methide and stabilizes the transition state that involves charge transfer to this moiety.

While many standalone pericyclases utilize binding energy to preorganize the linear substrate in a restricted NAC to overcome the rotational entropic barrier, LepI apparently does not function as “entropy trap” to increase reaction rate.^7,8,10,21,22,24,25^ The undehydrated substrate is bound in the extended conformation by LepI in order to minimize the π…π repulsive interaction between the diene and the electron-rich phenolate prior to dehydration. The slight entropy loss when the linear substrate rotates from extended conformation to the restricted NAC can be compensated by the gain of enthalpy from interaction between diene and electron-deficient quinone methide, and the enthalpy change that eventually occurs from the cycloaddition reaction. This may be a general feature for all dehydration-triggered pericyclases.

This feature is also reminiscent of another well-characterized pericyclase, chorismate mutase (CM), which catalyzes [3,3]-sigmatropic Claisen rearrangement of chorismate to prephenate. Unfavorable decreases in ΔS‡ have been consistently observed for natural CMs and catalytic antibodies, which suggests that CM does not act as entropy trap.^26^ Instead, the strategy of CM catalysis is a combination of both reactant destabilization and transition state stabilization.^26^ It has become clear that transition state stabilization is the more important contributor to CM catalysis: donating favorable hydrogen bond from cationic residue (such as Arg) to the vinyl ether oxygen in chorismate can effectively stabilize the developing partial negative charge of transition state.^27^ By replacing this critical arginine with its neutral analogue citrulline, which only confers hydrogen bond but not favorable electrostatic interactions, it is estimated that this positively charged arginine can contribute 5.9 kcal/mol to transition state stabilization versus 0.6 kcal/mol to the binding energy of the ground state.

Strikingly similar to CM, LepI uses similar cationic residues (H133 and R295) as major contributor for catalyzing [3,3]-sigmatropic retro-Claisen rearrangement of **9** to **10**. Substrate ground-state destabilization does not contribute, since the spirocyclic substrate **9** is essentially being frozen in the NAC like conformation, and no significant bond rotation is required to preorganize this substrate to be shape complementary to the LepI active site. The major contribution to catalysis must come from hydrogen bonding and electrostatic interaction. Substitution of H133 by neutral hydrogen bonding residue Gln and Asn reduces the catalytic activity by 50-fold. This severe loss of catalytic efficiency implies that H133 exists in the positively charged imidazolium form and underscores the importance of electrostatic complementarity. The importance of positively charged R295 to this pericyclic reaction has also been demonstrated by both QM studies and our current mutagenesis study.^12^. Consistent with this prediction, R295A/Q mutants are devoid of retro-Claisen rearrangement activity (>1,000 fold decrease, undetectable), whereas activity can be partially retained with similar cationic mutants: R295H (30%) (**Figure 6b**).

Although the precise catalytic role of SAM remains unclear, we have now found that the sulfonium does not contact the amide carbonyl. Nevertheless, the electrostatic effect of the sulfonium, or the ammonium analog can stabilize the transition state. Given that the positive charge of SAM is required for optimal activity of LepI, we propose that SAM serves as an electrostatic catalyst, stabilizing the transition state, which has a higher dipole moment than the reactant. This type of catalysis is well known through the work of Warshel, was demonstrated in solution chemistry by Wilcox, and has gained recent prominence under the name “electric field catalysis” through the work of Boxer, Shaik, Coote, and Head-Gordon.^28–33^

Our structural study of LepI also suggests an interesting evolutionary origin, that a methyltransferase ancestor of LepI has been coopted into a multifunctional dehydratase and pericyclase. The protein folds and SAM binding pockets are nearly identical for LepI and the methyltransferase, OxaC (**Supplementary Figure 7**). Comparison of the LepI active site with that of the OMT OxaC, along with structure-based sequence alignment (**Supplementary Figure 8**), reveals how key residues have been evolved to catalyze the reactions shown in Figure 1 instead of a SN2 methyl transfer reaction. In all functional *O*-MTs, a catalytic dyad His-Glu charge relay system is strictly conserved, in which the histidine acts as the general base to deprotonate the substrate nucleophile (hydroxyl group), and glutamate acts as the second shell residue to elevate the pKa of histidine. In LepI, the glutamate is conserved as E350, whereas the corresponding general base histidine is replaced with arginine. This substitution is beneficial for LepI in two ways. First of all, as discussed above, arginine is a better hydrogen bond catalyst and electrostatic catalyst in LepI as compared to histidine. Secondly, arginine is a poor general base due to higher pKa (12.5) and thus remains as protonated in the resting state. Removing this key general base in LepI may be a strategy to eliminate adventitious methyl-transfer activity.

In summary, our structural studies of LepI presented here, provide mechanistic insight of enzymatic dehydration triggered IMDA and HDA reactions, as well as hydrogen bonding and electrostatic catalysis of the retro-Claisen rearrangement. A proper understanding of the molecular mechanism of LepI is important for designing new pericyclases *de novo*, and also provides a template for the discovery of novel functions of OMT-like enzymes.

## Supporting information

## ACKNOWLEDGEMENT

This work was supported by the NSFC (9185620020) and the Strategic Priority Research Program (B) of CAS (XDB20000000) to JZ, NIH (1R01AI141481) and NSF (CHE-1806581) to YT and KNH. Chemical characterization studies were supported by shared instrumentation grants from the NSF (CHE-1048804) and the NIH NCRR (S10RR025631). We thank the staff of beamline BL17U1, BL18U1 and BL19U1 of Shanghai Synchrotron Radiation Facility (China) for access and help with the X-ray data collection. We also thank Prof. Jianhua Gan in Fudan University for help on structure refinement and Prof. Jiafu Long for help on ultracentrifugation sedimentation measurement. The computational resources from the UCLA Institute of Digital Research and Education (IDRE) are gratefully acknowledged. MO is supported by overseas postdoctoral fellowship from The Uehara Memorial Foundation, Japan. YH is a Life Sciences Research Foundation fellow sponsored by the Mark Foundation for Cancer Research.

## REFERENCES

1. Nicolaou KC, Snyder SA, Montagnon T, Vassilikogiannakis G (2002) The Diels—Alder reaction in total synthesis. Angew Chem Int Ed Engl 41:1668–1698.

2. Cooper SJ, Ohashi M, Liu F, Tang Y, Houk KN (2018) The expanding world of biosynthetic pericyclases: cooperation of experiment and theory for discovery. Nat Prod Rep Advance Article DOI:10.1039/c8np00075a

3. Chook YM, Ke H, Lipscomb WN (1993) Crystal structures of the monofunctional chorismite mutase from *Bacillus subtilis* and its complex with a transition state analog. Proc Natl Acad Sci USA 90:8600–8603.

4. Fage CD, Isiorho EA, Liu Y, Wagner DT, Liu HW, Keatinge-Clay AT (2015) The structure of SpnF, a standalone enzyme that catalyzes [4+2] cycloaddition. Nat Chem Biol 11:256–258.

5. Zaitseva J, Lu J, Olechoski KL, Lamb AL (2006) Two crystal structures of the isochorismate pyruvate lyase from *Pseudomonas aeruginosa*. J Biol Chem 281:33441–33449.

6. Shipman LW, Li D, Roessner CA, Scott AL, Sacchettini JC (2001) Crystal structure of precorrin-8x methyl mutase. Structure 9:587–596.

7. Zheng Q, et al. (2016) Enzyme-dependent [4+2] cycloaddition depends on lid-like interaction of the *N*-terminal sequence with the catalytic core in Pyrl4. Cell Chem Biol 23:352–360

8. Byrne MJ, et al. (2016) The catalytic mechanism of a natural Diels-Alderase revealed in molecular detail. J Am Chem Soc 138:6096–6098.

9. Cogan DP, et al. (2017) Structural insights into enzymatic [4+2] azacycloaddition in thiopeptide antibiotic biosynthesis. Proc Natl Acad Sci USA 114:12928–12933.

10. Zheng Q, et al. (2018) Structural insights into a flavin-dependent [4+2] cyclase that catalyzes *trans*-decalin formation in pyrroindomycin biosynthesis. Cell Chem Biol 25:718–727.

11. Newmister SA, et al. (2018) Structural basis of the Cope rearrangement and cyclization in hapalindole biogenesis. Nat Chem Biol 14:345–351.

12. Ohashi M. et al. (2017) SAM-dependent enzyme-catalysed pericyclic reactions in natural product biosynthesis. Nature 549:502–506.

13. Liscombe DK, Louie GV, Noel JP (2012) Architectures, mechanisms and molecular evolution of natural product methyltransferases. Nat Prod Rep 29:1238–1250.

14. Krissinel E, Henrick K (2007) Inference of macromolecular assemblies from crystalline state. J Mol Biol 372:774–797.

15. Newmister SA, et al. (2018) Unveiling sequential late-stage methyltransferase reactions in the meleagrin/oxaline biosynthetic pathway. Org Biolmol Chem 16:6450–6459.

16. Holm L, Rosenström P (2010) Dali server: conservation mapping in 3D. Nucleic Acids Res 38:w545–549.

17. Horowitz S, et al. (2013) Conservation and functional importance of carbon-oxygen hydrogen bonding in AdoMet-dependent methyltransferases. J Am Chem Soc 135:15536–15548.

18. Fick RJ, et al. (2018) Water-mediated carbon-oxygen hydrogen bonding facilitates S-adenosylmethionine recognition in the reactivation domain of cobalamin-dependent methionine synthase. Biochemistry 57:3733–3740

19. Yu J, Zhou Y, Tanaka I, Yao M (2010) Roll: a new algorithm for the detection of protein pockets and cavities with a rolling probe sphere. Bioinformatics 26:46–52.

20. Pauling L (1946) Molecular architecture and biological reactions. Chem Eng News 24:1375–1377

21. Jeon BS, Wang SA, Ruszczycky MW, Liu HW (2017) Natural [4+2]-cyclases. Chem Rev 117:5367-5388

22. Preiswerk N, et al. (2014) Impact of scaffold rigidity on the design and evolution of an artificial Diels-Alderase. Proc Natl Acad Sci USA 111:8013–8018.

23. Minami A, Oikawa H (2016) Recent advances of Diels-Alderases involved in natural product biosynthesis. J Antibiot 69:500–506.

24. Breslow R (1991) Hydrophobic effects on simple organic reactions in water. Acc Chem Res 24:159–164.

25. Blokzijl W, Blandamer MJ, Engberts JBFN (1991) Diels-Alder reactions in aqueous solutions. Enforced hydrophobic interactions between diene and dienophile. J Am Chem Soc 113:4241–4246.

26. Lee AY, Stewart JD, Clardy J, Ganem B (1995) New insight into the catalytic mechanism of chorismite mutase from structural studies. Chem Biol 2:195–203.

27. Burschowsky D, et al. (2014) Electrostatic transition state stabilization rather than reactant destabilization provides the chemical basis for efficient chorismite mutase catalysis. Proc Natl Acad Sci USA 111:17516–17521.

28. Warshel A, et al. (2006) Electrostatic Basis for Enzyme Catalysis. Chem Rev 106:3210–3235.

29. Smith PJ, Wilcox CS (1991) The chemistry of functional group arrays. Electrostatic catalysis and the “intramolecular salt effect”. Tetrahedron 47:2617–2628.

30. Fried SD, Boxer SG (2017) Electric Fields and Enzyme Catalysis. Annu Rev Biochem 86:387–415.

31. Shaik S, Mandal D, Ramanan, R (2016) Oriented electric fields as future smart reagents in chemistry. Nature Chem 8:1091–1098.

32. Aragonès AC, et al. (2016) Electrostatic catalysis of a Diels-Alder reaction. Nature 531:88–91.

33. Welborn VV, Ruiz PL, Head-G T (2018) Computational optimization of electric fields for better catalysis design. Nature Catal 1:649–655.

