## Supplementary material for "Structural Basis for Stereoselective Dehydration and Hydrogen-Bonding Catalysis by the SAM-Dependent Pericyclase LepI"

#### METHODS

##### General DNA manipulation technique

The wild-type LepI gene was subcloned into a pET-28a derivative vector to express LepI as an N-terminal $(His)_6$ tag fused protein. Mutations were introduced by PCR into the above plasmid using the QuikChange kit (Stratagene). The gene-specific primers are listed in Supplementary Table 2. The sequence of each construct was confirmed by DNA sequencing.

##### Protein Expression and Purification.

The wild-type LepI protein was expressed in *E. coli* BL21 (DE3) in LB medium in the presence of 50 mg/L kanamycin. Expression was induced by 0.4 mM IPTG (isopropyl-β-D-thiogalactopyranoside) when $OD_{600}$ reached 1.0. The selenomehionine-derivatized (Se-Met) LepI was overexpressed in *E. coli* B834 (DE3) in M9 medium at 37 °C in the presence of 50 mg/L kanamycin until $OD_{600}$ reached 0.6. After supplementation with 50 mg/L L-(+)-selenomethionine (J&K Scientific), Se-Met protein expression was induced by the addition of 0.4 mM IPTG. After growing for 16 h at 16°C, the cells were harvested, homogenized in a buffer containing 25 mM tris (pH 8.0), 300 mM NaCl, 5 mM β-mercaptoethanol, 1 mM phenylmethylsulfonyl fluoride (PMSF), and lysed by French press with a high-pressure homogenizer (60-100 MPa). Cell debris was removed by centrifugation at 16000 rpm for 30 min at 4°C. The supernatant was loaded onto a Ni-NTA affinity column (GE), and the His-tagged protein was collected by elution with a buffer containing 25 mM Tris (pH 8.0), 300 mM NaCl, 300 mM imidazole, 5 mM 2-mercaptoethanol, 10% glycerol) and further concentrated using Amicon ultra filter units (Millipore) with a 30 kDa molecular weight cutoff. The concentrated protein was then applied to a HiLoad Superdex 200 column (GE Healthcare) in a buffer containing 25 mM tris (pH 8.0), 150 mM NaCl and 3 mM DTT. The peak fractions were collected and concentrated for crystallization and activity assay. A surface mutant C52A was expressed and purified similarly. This mutant was prepared during our crystallization optimization process, and was found to substantially facilitate cocrystallization and improve the quality of cocrystals. Mutants of LepI were purified by the same method. A composite of all purified LepI and mutants can be found in **Supplementary Figure 8**.

**Protein Crystallization**

Crystals of the Se-Met LepI were grown at 16 °C using the sitting drop vapor diffusion method in 2 μL drops containing an 1:1 mixture of the protein solution (20 mg/mL Se-Met LepI in buffer containing 25 mM tris (pH8.0), 150 mM NaCl and 3 mM DTT) and a reservoir solution consisting of 0.1 M MES/NaOH (pH 6.0), 20% PEG 6000 and 200 mM $AlCl_3$. ). Prism-like crystals appeared after one day at 16 °C. Crystals of native LepI were grown at 16 °C using the sitting drop vapor diffusion method in 2 μL drops containing an 1:1 mixture of the protein solution (20 mg/mL LepI in buffer containing 25 mM tris (pH8.0), 150 mM NaCl and 3 mM DTT) and a reservoir solution consisting of 0.1 M MES/NaOH (pH 6.0), 20% PEG 6000 and 200 mM NaCl. Prism-like crystals appeared after 1 day at 16 °C. No exogenous SAM was added in protein expression, purification and crystallization.

All LepI-SAM-compound tertiary complexes were crystallized using the sitting drop vapor diffusion method at 16 °C. Purified LepI (20 mg/mL) was first incubated with 2.5 mM compound and 3-5% DMSO in buffer containing 25 mM tris (pH8.0), 150 mM NaCl and 3 mM DTT on ice for 30 min and followed by centrifuge. 1 μL of protein solution was mixed with 1 μL of precipitant solution. Prism-like crystals appeared after 1 day at 16 °C. The LepI-SAM-**1** complex crystals were grown in 0.1 M ADA (pH 6.5), 12% PEG 6000 and 100 mM $MgCl_2$). The

C52A-SAH-**1** was grown under the same condition. The LepI-SAM-**8** complex crystals were grown in 0.1 M MES/NaOH (pH 6.0), 20% PEG 6000 and 200 mM $AlCl_3$. The LepI-SAM-**10** complex was crystallized in 0.1 M sodium citrate/citric acid (pH 5.5), 15% PEG 10000 and 2% dioxane. No exogenous SAM or SAH was added in protein crystallization.

All crystals were flash-frozen in liquid nitrogen after being transferred to a cryoprotectant solution consisting of mother liquor supplemented with 10-15% (v/v) glycerol.

**Data Collection and Structure Determination**

All X-ray diffraction data were recorded at the Shanghai Synchrotron Radiation Facility (SSRF). For Se-Met LepI, data were collected at BL19U1 ($\lambda$ = 0.97855 Å). For native LepI, LepI-SAM-**1**, LepI-SAM-**10** and LepI-SAM-**8** complex, data were collected at BL18U1 ($\lambda$ = 0.97930 Å). For C52A-SAM-ketone complex, data were collected at BL17U1 ($\lambda$ = 0.97915 Å). Data reduction and integration for C52A-SAM-ketone data sets was achieved with HKL2000 package[1] while others were achieved with HKL3000 package.[2] The statistics for data collection are listed in **Supplementary Table 1**. All crystals belonged to space group *C2* with two molecules in the asymmetric unit.

Structure of SeMet-LepI in complex with SAM was determined by SAD. 27 selenium sites were identified in one asymmetric unit by Autosol in the PHENIX package and used for model building through Autobuild.[3] Structures of LepI-SAM complex were determined by molecular replacement using Phaser[4] and the atomic coordinates of SeMet-LepI-SAM complex as the search model. Iterative cycles of model rebuilding and refinement were carried out using COOT[5], Refmac5[6], and PHENIX[3]. TLSNative LepI structure was determined by molecular replacement using the program PHASER and the atomic coordinates of one chain with the highest completion after Autobuild as the search model.[4] Structures of other LepI complexes were determined by molecular replacement using PHASER[4] and the atomic coordinates of LepI-SAM complex as the search model. PROCHECK[7] and MolProbity[8] were used to access the overall quality of the structural models. Refinement statistics for each final model are recorded in **Supplementary Table 1**. Structure figures were made using PyMol 1.3 (Schrödinger, LLC).[9]

**Construction of plasmids for expression of LepI mutants.**

The oligonucleotide primers used for generating the mutants are listed in **Supplementary Table 2**. The plasmid pMO0048[10] containing the wild-type *lepI* gene was used as the template for PCR-based site-directed mutagenesis. The primers of LepI-M-f1/D296N-r1 and D296N-f1/LepI-M-r1 were used for Asp296Asn, and the resulting two overlapped fragments and the pET28b(+) expression vector amplified using the primers of LepI-vec-f1/r1 were combined to generate the Asp296Asn plasmid using GeneArt Seamless Cloning and Assembly kit (Thermo Fisher Scientific). Other mutants were constructed in the same manner using primer pairs (ex. LepI-M-f1/mutation position-r1 and mutation position-f1/LepI-M-r1). DNA sequencing was used to confirm the identities including the mutated positions of the expression plasmids.

***In vitro* analysis of the retro-Claisen activity of LepI mutants using compound 9 as the substrate**

The procedure was as described previously.[10] Assays for the activity of LepI mutants with **9** in HEPES buffer (50 mM HEPES, 100 mM NaCl, pH 8.0) were performed at 50 μL scale with LepI mutants at 30 °C for 5 min. Then the reaction was quenched with equal volume of cold acetonitrile. Protein was precipitated and removed by centrifugation and the supernatant analyzed by HPLC using a C18 column (Phenomenex Luna C18 (2) 5 μm, 2.0 × 100 mm) with isoclatic conditions (50% of $H_2O$ in $CH_3CN$). Results were quantified by the standard curve of product **10**. Final results were calculated as percent of controls. The error bars represent standard deviation (s.d.) of three independent replicates. These results are shown in **Figure 6b**.

**Preparation of overexpression plasmids of lepI for *A. nidulans***

pMO0008, pMO0011, and pMO0035 were prepared as described previously.[10] The gpdA promoter and trpC terminator were amplified with primers pMO0107-f1/r1 and pMO0107-f3/r3, respectively. The gene of lepI and the lepI mutants were amplified using the corresponding protein expression vectors as the template with primers pMO0107-f2/r2. The three overlapping DNA fragments and PshAI/NotI-digested pMO0011 expression vector were transformed into yeast to generate the desired plasmids by yeast homologous recombination. DNA sequencing was used to confirm the identities of the expression plasmids. The plasmids used for the heterologous expression are illustrated in **Supplementary Figure 10**.

**Heterologous expression of *lep* genes in *A. nidulans* and the analysis of metabolites**

The procedure is as described previously.[10] *A. nidulans* A1145 was initially grown on CD agar plates containing 10 mM uracil, 0.5 μg/mL pyridoxine·HCl, and 2.5 μg/mL riboflavin at 30 °C for 5 days. Fresh spores of *A. nidulans* were inoculated into 40 mL liquid CD media (1 L: 10 g Glucose, 50 mL 20 x Nitrate salts, 1 mL Trace elements, pH 6.5) in 250-mL Erlenmeyer flask and germinated at 30 °C with shaking at 250 rpm for approximately 16 h. Mycelia were harvested by centrifugation at 3,500 rpm for 10 min and washed with 10 mL osmotic buffer (1.2 M $MgSO_4 \cdot 7H_2O$, 10 mM sodium phosphate, pH 5.8). Then the mycelia were suspended in 10 mL of osmotic buffer with 30 mg lysing enzymes from *Trichoderma* and 20 mg Yatalase and transferred to a 125-mL flask. The flask was shaken at 80 rpm for overnight at 30 °C. Cells were collected in a 30 mL Corex tube and overlaid gently with 10 mL of Trapping buffer (0.6 M sorbitol, 0.1 M Tris-HCI, pH 7.0). After centrifugation at 3,500 rpm for 15 min at 4 °C, protoplasts were collected from the interface of the two buffers, transferred to a sterile 15-mL falcon tube and washed with 10 mL STC buffer (1.2 M sorbitol, 10 mM $CaCl_2$, 10 mM Tris-HCI, pH 7.5). The protoplasts were resuspended in 1 mL STC buffer for transformation. Then, the plasmids (see Supplementary Information) were added to 100 μL protoplast suspension. After incubating the mixture on ice for 60 min, 600 μL of PEG solution at pH 7.5 (60% PEG, 50 mM $CaCl_2$ and 50 mM Tris-HCl) was added to the protoplast mixture, and the mixture was incubated at room temperature for additional 20 min. The mixture was spread on the regeneration dropout solid medium (CD solid medium with 1.2 M sorbitol and appropriate supplements) and incubated at 30 °C for 3 days.

For analysis, the transformants of *A. nidulans* strains were grown for 3 days in 20 mL liquid CD-ST. 500 μL of cells were collected and extracted with 750 μL EtOAc/MeOH mixtute (10:1, v/v). The organic phase was dried by speed vacuum and dissolved in MeOH for analysis. LC–MS analyses were performed on a Shimadzu 2020 EV LC–MS (Kinetex 1.7 μm C18 100 Å, LC Column 100 × 2.1 mm) using positive-and negative-mode electrospray ionization with a linear gradient of 5–95% MeCN–$H_2O$ with 0.5% formic acid in 15 min followed by 95% MeCN for 3 min with a flow rate of 0.3 mL/min. The results are shown in **Figure 6a**.

**Isolation and structural characterization of compound 8 from the transformant of *A. nidulans***

For isolation of the IMDA adduct **8**, the transformants of *A. nidulans* strains harboring pMO0008 (LepA), pMO0011 (LepH), and pMO0035 (LepG and LepF) were grown for 84 h in 4 x 1 L liquid CD-ST and filtered to collect the cells from liquid culture. The cells were extracted with 1 L acetone, and the extracts were evaporated to dryness and partitioned twice between EtOAc/$H_2O$. After evaporation of the organic phase, the crude extracts were separated by silica gel chromatography with *n*-hexane/EtOAc (1:0 to 0:1). Fractions containing the IMDA adduct **8** were combined and purified by HPLC with a semi-preparative C18 column of Kinetics New column, 5 μm, 10 × 250 mm using an isocratic elution system of 55 % MeCN (v/v) in $H_2O$ with 0.05% (v/v) formic acid at a flow rate of 4.0 mL/min to afford the mixture of **8** and **5 (**20 mg**)**. The mixture was used as a ligand for the co-crystalization with LepI. As a result, LepI-SAM-**8** tertiary complexes were obtained. Since **8** was not characterized in the previous study, to separate **8** and **5**, the mixture was further purified by HPLC with an analytical chiral column of Lux Cellulose-1 column, 5 μm, 4.6 × 250 mm using an isocratic elution system of 65 % MeCN (v/v) in $H_2O$ with 0.05% (v/v) formic acid at a flow rate of 1.0 mL/min to afford pure **8** (3.0 mg). For elucidation of chemical structure of **8**, 1D and 2D NMR spectra were obtained on Bruker AV500 spectrometer at the UCLA Molecular Instrumentation Center and are shown in **Supplementary Figures 11-15**. High-resolution mass spectra were obtained from Thermo Fisher Scientific Exactive Plus with IonSense ID-CUBE DART source at the UCLA Molecular Instrumentation Center. To determine the streochemistry of **8**, a single crystal of **8** was prepared from MeCN/$H_2O$.

**Computational Methods**

The DFT calculations were performed with Gaussian 09.[11] Geometry optimizations of all the minima and transition state structures were carried out at the B3LYP-D3/6-31G(d) level of theory.[12-16] This level of theory was previously benchmarked against M06-2X and ωB97X-D and shown to yield comparable results.[10,17] Docking files were prepared using AutoDock Tools 1.5.6.[18] Structures were docked into the active site of LepI using AutoDock Vina with an exhaustiveness of 12 and a 30 Å grid cube (27,000 Å$^3$).[19]

**Ultracentrifugation analysis of LepI**

Sedimentation velocity experiments were performed on a Beckman Optima AUC analytical ultracentrifuge equipped with an eight-cell rotor under 35,000 rpm at 4°C. The partial specific volume of different protein samples and the buffer density were calculated using the program SEDNTERP (http://www.rasmb.bbri.org/). The final sedimentation velocity data were analyzed and fitted to a continuous sedimentation coefficient distribution model using the program SEDFIT.[20]

**Supplemental References**

1. Otwinowski Z, Minor W (1997) Processing of X-ray diffraction data collected in oscillation mode. *Methods Enzymol* 276:307–326.

2. Minor W, Cymboriwski M, Otwinowski Z, Chruszcz M (2006) HKL-3000: the integration of data reduction and structure solution-from diffraction images to an initial model in minutes. *Acta Crystallogr D Biol Crystallogr* 62:859–866.

3. Adams PD, et al. (2010) PHENIX: A comprehensive Python-based system for macromolecular structure solution. *Acta Crystallogr* D66:213−221.

4. McCoy AJ, et al. (2007) Phaser crystallographic software. *J Appl Cryst* 40:658–674.

5. Emsley P, Lohkamp B, Scott WG, Cowtan K (2010) Features and development of Coot. *Acta Crystallogr* D66:486−501.

6. Murshudov GN, Vagin AA, Dodson EJ (1997) Refinement of macromolecular structures by maximum-likelihood method. *Acta Crystallogr D Biol Crystallogr* 53:240–255.

7. Laskowski RA, MacArthur MW, Moss DS, Thornton JM (1993) ProCheck: a program to check the stereochemical quality of protein structures. *J Appl Cryst* 26:283–291.

8. Davis I, W. et al. (2007) MolProbity: all-atom contacts and structure validation for proteins and nucleic acids. *Nucleic Acids Res* 35(Web Server issue):W375–W383.

9. DeLano WL (2002) PyMOL: an open-source molecular graphics tool. *Ccp4 Newslett Protein Crystallogr* 40:11.

10. Ohashi M, et al. (2017) SAM-dependent enzyme-catalysed pericyclic reactions in natural product biosynthesis. *Nature* 549:502–506.

11. Frisch MJ, et al. (2013) Gaussian 09 (Gaussian, Inc., Wallingford CT).

12. Becke AD (1993) Density-functional thermochemistry. III. The role of exact exchange. *J Chem Phys* 98:5648–5652.

13. Lee C, Yang W, Parr RG (1988) Development of the Colle–Salvetti correlation-energy formula into a functional of the electron density. *Phys Rev B* 37:785–789.

14. Vosko SH, Wilk L, Nusair M (1980) Accurate spin-dependent electron liquid correlation energies for local spin density calculations: A critical analysis. *Can J Phys* 58:1200–1211.

15. Stephens PJ, Devlin FJ, Chabalowski CF, Frisch MJ (1994) *J Phys Chem* 98:11623–11627.

16. Grimme S, Antony J, Ehrlich S, Krieg H (2010) A consistent and accurate ab initio parametrization of density functional dispersion correction (DFT-D) for the 94 elements H-Pu. *J Chem Phys* 132:154104.

17. Patel A, et al. (2016) Dynamically complex [6+4] and [4+2] cycloadditions in the biosynthesis of spinosyn A. *J Am Chem Soc* 138:3631–3634.

18. Morris GM, et al. (2009) Autodock4 and AutoDockTools4: automated docking with selective receptor flexiblity. *J. Computational Chemistry* 16:2785–2791.

19. Trott O, Olson AJ (2010) AutoDock Vina: Improving the speed and accuracy of docking with a new scoring function, efficient optimization, and multithreading. *J Comput Chem* 31:455–461.

20. Schuck, P. (2000) Size-distribution analysis of macromolecules by sedimentation velocity ultracentrifugation and lamm equation modeling. *Biophys J* 78:1606–1619.

### SUPPLEMENTARY TABLES

#### Supplementary Table 1 | Data collection and refinement statistics.

| | LepI-SAM complex | LepI-SAM-**1** complex | LepI(C52A)-SAH-**1** complex | LepI(C52A)-SAM-**10** complex | LepI(C52A)-SAM-**8** complex |
|---|---|---|---|---|---|
| **Data collection** | | | | | |
| Space group | *C2* | *C2* | *C2* | *C2* | *C2* |
| Cell dimensions | | | | | |
| a, b, c (Å) | 161.3, 61.8, 113.3 | 161.1, 62.2, 114.2 | 161.3, 62.1, 113.6 | 161.3, 62.3, 113.9 | 160.8, 62.6, 113.8 |
| α, β, γ (°) | 90.0, 113.5, 90.0 | 90.0, 113.1, 90.0 | 90.0, 113.7, 90.0 | 90.0, 113.5, 90.0 | 90.0, 113.3, 90.0 |
| Resolution (Å) | 50.00 - 2.14 (2.18 - 2.14) | 50.00 - 1.70 (1.73 - 1.70) | 50.00 - 1.83 (1.86 - 1.83) | 50.00 - 1.78 (1.81 - 1.78) | 50.00 - 1.66 (1.69 - 1.66) |
| $R_{merge}$ | 0.192 (2.028) | 0.057 (0.876) | 0.101 (1.647) | 0.095 (1.932) | 0.066 (1.072) |
| $R_{pim}$ | 0.066 (0.750) | 0.024 (0.368) | 0.035 (0.555) | 0.027 (0.616) | 0.028 (0.463) |
| $I/\sigma I$ | 19.6 (4.1) | 30.7 (2.2) | 35.0 (7.5) | 33.6 (1.8) | 25.8 (2.0) |
| Completeness (%) | 97.9 (97.2) | 99.9 (99.9) | 98.7 (99.8) | 99.8 (99.3) | 99.7 (99.0) |
| Redundancy | 9.1 (7.8) | 6.6 (6.5) | 9.1 (9.6) | 13.0 (10.1) | 6.2 (6.2) |
| $CC_{1/2}$ | 0.984 (0.787) | 0.994 (0.809) | 0.996 (0.920) | 1.002 (0.692) | 0.997 (0.731) |
| **Refinement** | | | | | |
| Resolution (Å) | 36.96 - 2.13 (2.21 - 2.13) | 34.99 - 1.70 (1.76 - 1.70) | (32.96 - 1.84) (1.90 - 1.84) | 35.96 - 1.78 (1.84 - 1.78) | 28.80 - 1.66 (1.72 - 1.66) |
| No. reflections | 55671 (4947) | 111704 (9075) | 89085 (8908) | 95303 (5881) | 117502 (8551) |
| $R_{work}$ / $R_{free}$ | 0.184/0.218 (0.215/0.269) | 0.163/0.189 (0.224/0.260) | 0.138/0.173 (0.128/0.190) | 0.185/0.208 (0.268/0.302) | 0.168/0.197 (0.231/0.272) |
| No. atoms | | | | | |
| Protein | 6232 | 6409 | 6285 | 6466 | 6342 |
| Ligand | 55 | 137 | 90 | 33 | 77 |
| Water | 452 | 929 | 680 | 550 | 692 |
| *B* factors ($Å^2$) | | | | | |
| Protein | 27.6 | 16.3 | 21.6 | 27.1 | 17.2 |
| Ligand/ion | 26.4 | 30.4 | 30.8 | 34.6 | 27.3 |
| Water | 36.0 | 29.1 | 33.4 | 34.1 | 26.7 |
| R.m.s. deviations | | | | | |
| bond lengths (Å) | 0.003 | 0.014 | 0.007 | 0.010 | 0.018 |
| bond angles (°) | 0.77 | 1.3 | 0.9 | 1.1 | 1.5 |
| Ramachandran outliers | 0.00% | 0.00% | 0.00% | 0.13% | 0.13% |
| Ramachandran favored | 97.67% | 97.93% | 97.93% | 97.42% | 98.06% |
| **PDB code** | **6IX3** | **6IX5** | **6IX7** | **6IX9** | **6IX8** |

**Supplementary Table 2.** Primers used in this study

| Primer | Sequences of primer (5'→3') |
|---|---|
| LepI-M-f1 | TTTAACTTTAAGAAGGAGATATACCATGGAGACTGTCGCTGCCATAAAG |
| LepI-M-r1 | CAACTCAGCTTCCTTTCGGGCTTTG |
| LepI-vec-f1 | CAAAGCCCGAAAGGAAGCTG |
| LepI-vec-r1 | GGTATATCTCCTTCTTAAAGTTAAACAAAATTATTTC |
| D296N-f1 | GTCTCATTCTGCGCAACTTCCCTGA |
| D296N-r1 | TCAGGGAAGTTGCGCAGAATGAGAC |
| D296A-f1 | GTCTCATTCTGCGCGCGTTCCCTGACC |
| D296A-r1 | GGTCAGGGAACGCGCGCAGAATGAGAC |
| D296E-f1 | GTCTCATTCTGCGCGAATTCCCTGACC |
| D296E-r1 | GGTCAGGGAATTCGCGCAGAATGAGAC |
| R295K-f1 | CCGTCTCATTCTGAAAGACTTCCCTGACC |
| R295K-r1 | GGTCAGGGAAGTCTTTCAGAATGAGACGG |
| R295E-f1 | CCGTCTCATTCTGGAAGACTTCCCTGACCA |
| R295E-r1 | TGGTCAGGGAAGTCTTCCAGAATGAGACGG |
| R295A-f1 | GGTCAGGGAAGTCCGCCAGAATGAGACGG |
| R295A-r1 | AGCAGGAATGCGAGCGGCATTCGAGAATC |
| R295H-f1 | GTCTCATTCTGCATGACTTCCCTGA |
| R295H-r1 | TCAGGGAAGTCATGCAGAATGAGAC |
| R295Q-f1 | CCGTCTCATTCTGCAGGACTTCCCTGACCAC |
| R295Q-r1 | GTGGTCAGGGAAGTCCTGCAGAATGAGACGG |
| R295N-f1 | CCGTCTCATTCTGAACGACTTCCCTGACCAC |
| R295N-r1 | GTGGTCAGGGAAGTCGTTCAGAATGAGACGG |
| R295F-f1 | CCGTCTCATTCTGTTTGACTTCCCTGACC |
| R295F-r1 | GGTCAGGGAAGTCAAACAGAATGAGACGG |
| R295Y-f1 | CCGTCTCATTCTGTATGACTTCCCTGACC |
| R295Y-r1 | GGTCAGGGAAGTCATACAGAATGAGACGG |
| H133A-f1 | GGAATGCGAGCGGCATTCGAGAATC |
| H133A-r1 | GATTCTCGAATGCCGCTCGCATTCC |
| H133F-f1 | GCAGGAATGCGATTTGCATTCGAGAATC |
| H133F-r1 | GATTCTCGAATGCAAATCGCATTCCTGC |
| R197A-f1 | CTCACTGACGAGGCGACCCCAAACTTC |
| R197A-r1 | GAAGTTTGGGGTCGCCTCGTCAGTGAG |
| R197K-f1 | CTCACTGACGAGAAAACCCCAAACTTCC |
| R197K-r1 | GGAAGTTTGGGGTTTTCTCGTCAGTGAG |
| T338A-f1 | GCAGAAACCGGAGCGGACATCTGCATT |
| T338A-r1 | AATGCAGATGTCCGCTCCGGTTTCTGC |
| T338S-f1 | GGCAGAAACCGGATCTGACATCTGCAT |
| T338S-r1 | ATGCAGATGTCAGATCCGGTTTCTGCC |
| M45A-f1 | CGATACGGTGGCGCGGATGTCGCTC |
| M45A-r1 | GAGCGACATCCGCGCCACCGTATCG |
| pMO0107-f1 | CGCGGGTGTTCTTGACGATGGCATCCTGCGGCCGCACTCCGGTGAATTGATTTGGGTG |
| pMO0107-r1 | TGTTTAGATGTGTCTATGTGGCGG |
| pMO0107-f2 | TAACCATTACCCCGCCACATAGACACATCTAAACAATGGAGACTGTCGCTGCCATAA AG |
| pMO0107-r2 | TCAAGCTGTTTGATGATTTCAGTAACGTTAAGTGGTCACTGCAGGCTAAACACCATG |
| pMO0107-f3 | CCACTTAACGTTACTGAAATCATCAAAC |
| pMO0107-r3 | TCATTTATAGCTCGTTCGGCACCTTTAATCAAGAAGGATTACCTCTAAACAAGTGTAC CT |

**Supplementary Table 3. Spectroscopic data of compound 8.**

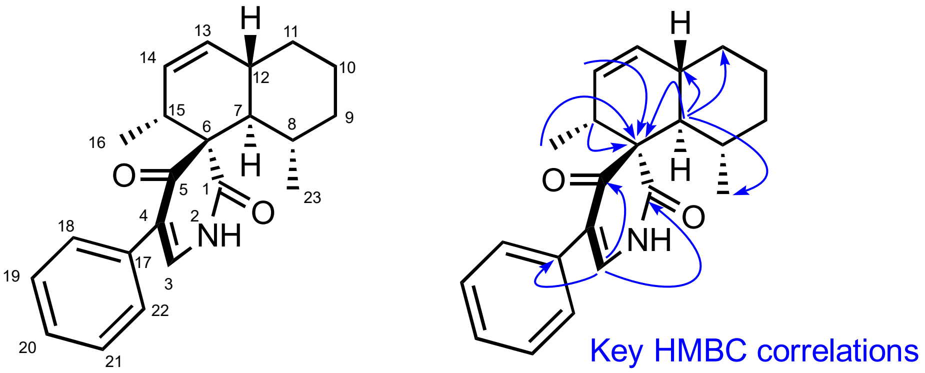


| position | $\delta_H$, mult (*J* in Hz) | $\delta_C$ | COSY |
|---|---|---|---|
| 1 | | 178.04 | |
| 2 | 8.51, brs | | H3 |
| 3 | 7.33, overlapped | 138.96 | H2 |
| 4 | | 120.51 | |
| 5 | | 197.61 | |
| 6 | | 65.34 | |
| 7 | 1.90, dd (11.0, 11.0) | 50.8 | H8, H12 |
| 8 | 1.76, m | 34.05 | H7, H9, H23 |
| 9 | 1.67, m<br>1.04, m | 36.44 | H8, H10 |
| 10 | 1.69, m<br>1.39, m | 25.74 | H9, H11 |
| 11 | 2.03, m<br>1.24, m | 33.77 | H10, H12 |
| 12 | 2.48, m | 38.48 | H7, H11, H13, H14 |
| 13 | 5.85-5.80, m | 137.37 | H14, H12 |
| 14 | 5.56, ddd (9.5, 3.0, 3.0) | 128.52 | H13, H15 |
| 15 | 2.82-2.70, m | 41.97 | H13, H14, H16 |
| 16 | 1.13, d (7.5) | 16.77 | H15 |
| 17 | | 133.42 | |
| 18/22<br>19/21<br>20 | 7.40-7.29, m | 128.62<br>128.52<br>127.90 | |
| 23 | 0.75, d (6.5) | 21.17 | H8 |

$^1$H NMR (500 MHz) and $^{13}$C NMR (125 MHz) data of compound **8** in $CDCl_3$. The absolute configuration is shown. The absolute stereochemistry of **8** was determined by single crystal X-ray analysis (**Supplementary material**) and LepI-SAM-compound **8** tertiary complexes (**Figure 5**). HRMS (ESI, $MH^+$) calcd for $C_{22}H_{26}NO_2$ 336.1958, found 336.1943.

#### SUPPLEMENTARY FIGURES

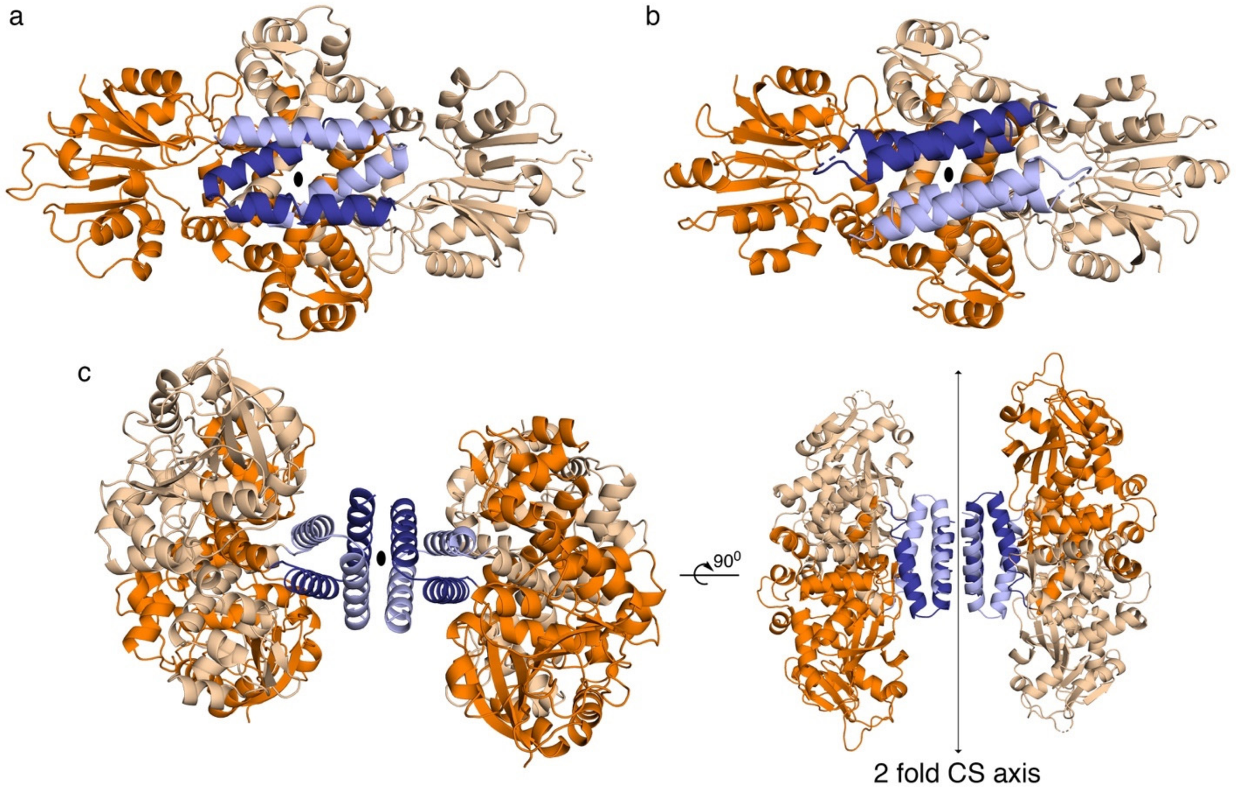


**Supplementary Figure 1 | Quaternary structure of LepI and comparison with *O*-methyltransferase OxaC.** (a) Top view of LepI homodimer. Protomers are colored in dark and pale color, respectively. The *N*-terminal domain-swapped coiled-coil helical bundle is shown in blue. The noncrystallographic two-fold rotational axis running perpendicular through the center. (b) Top view of OxaC homodimer, color coded similar to LepI, highlighting different fold of the *N*-terminal helical bundle. The crystallographic two-fold axis is shown. (c) Top and side view of LepI tetramer (a dimer of dimers). The crystallographic 2-fold rotational axis is shown at the interface.

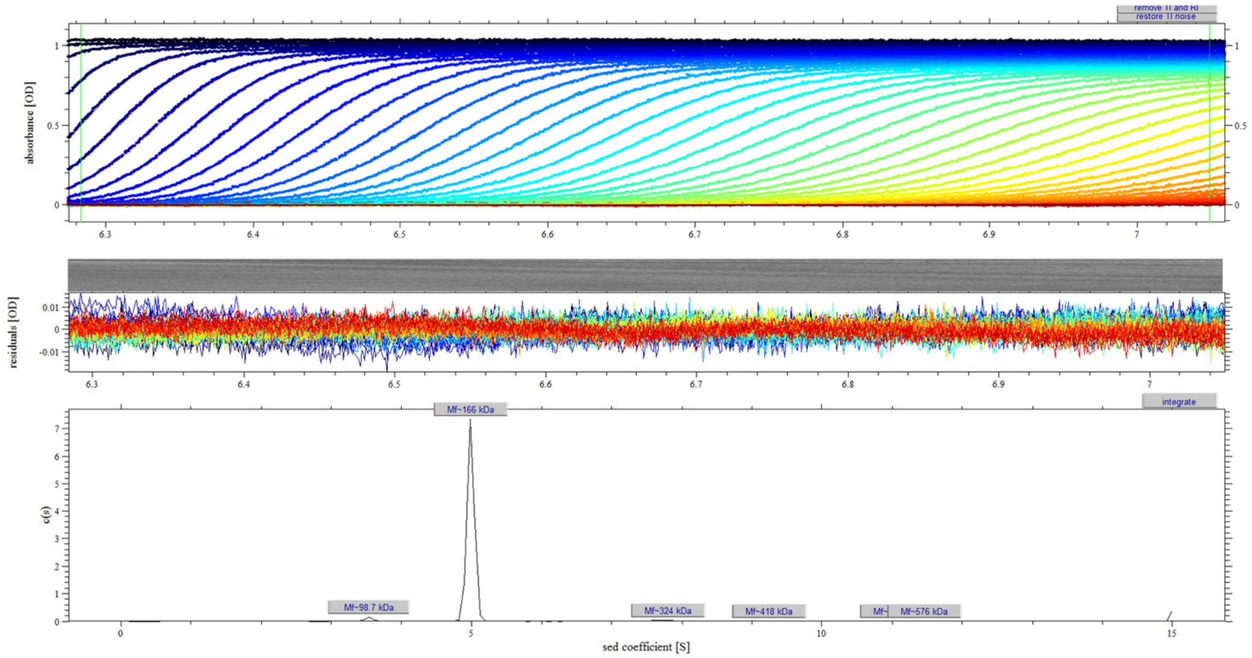


**Supplementary Figure 2 | Solution oligomerization state of LepI characterized by ultracentrifugation velocity.** The major peak has a sedimentation coefficient consistent with a LepI tetramer while 2% of dimer also existed in solution.

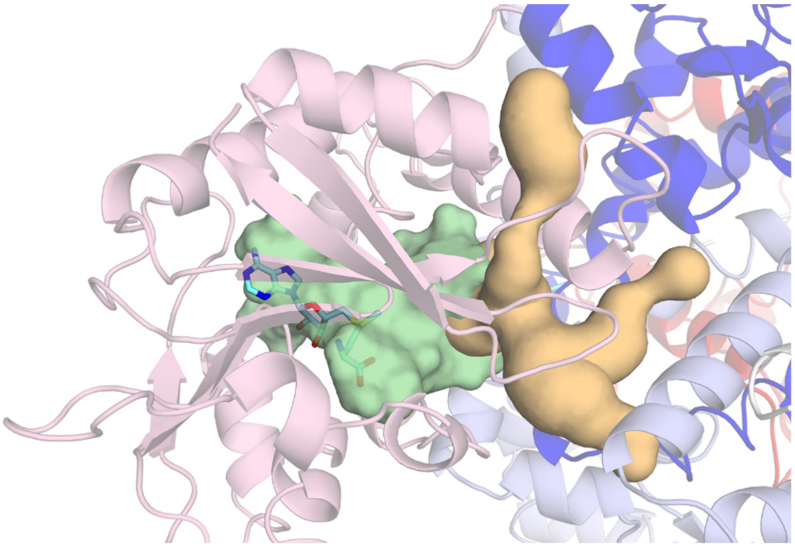

**Supplementary Figure 3 | Solvent-releasing tunnels at the domain interface.** The tunnels are shown in yellow surface**.** The active site cavity including both SAM binding site and substrate binding site is shown in green surface. SAM is shown in stick model.

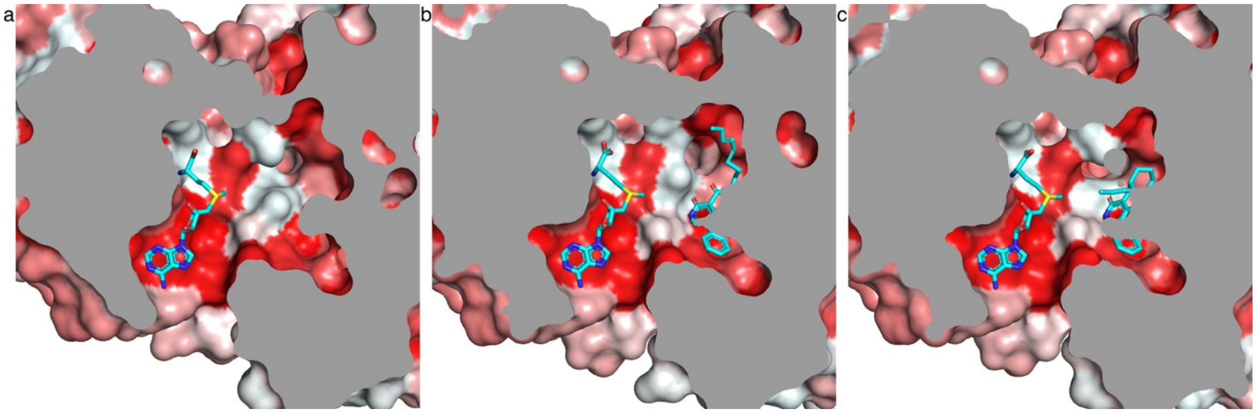


**Supplementary Figure 4 | Clip view of active site cavity.** The hydrophobic surface is shown in red, whereas the hydrophilic surface is shown in white. (a) ligand-free active site. (b) linear substrate analogue ketone **1** in the active site. (c) cyclic-product leporin C **10** in the active site.

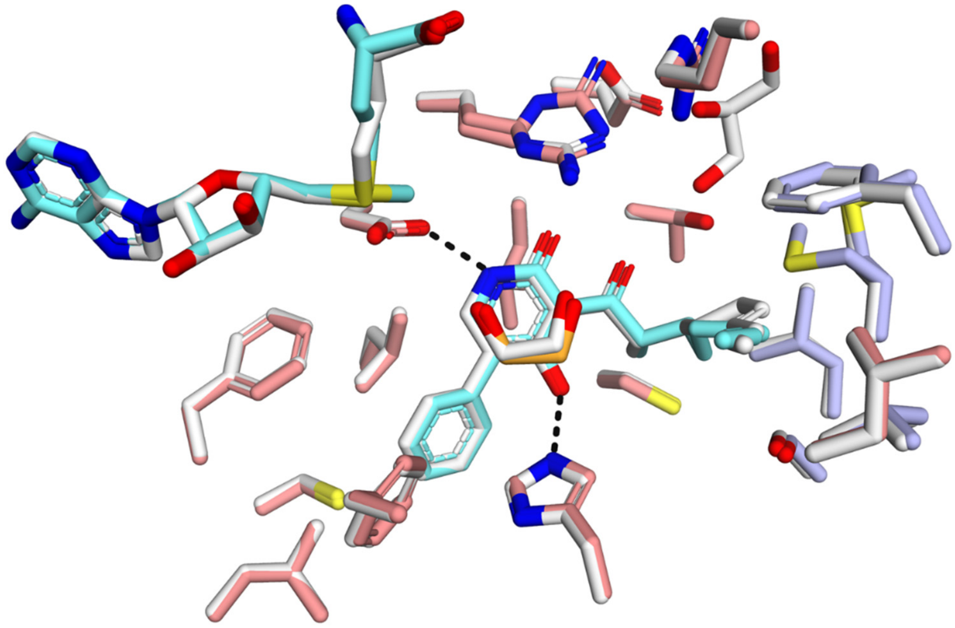

**Supplementary Figure 5 | Structural comparison between LepI-SAM-1 and LepI-SAH-1 ternary complex.** LepI-SAM-**1** ternary is color-coded as in Figure 3. LepI-SAH-1 is colored in white.

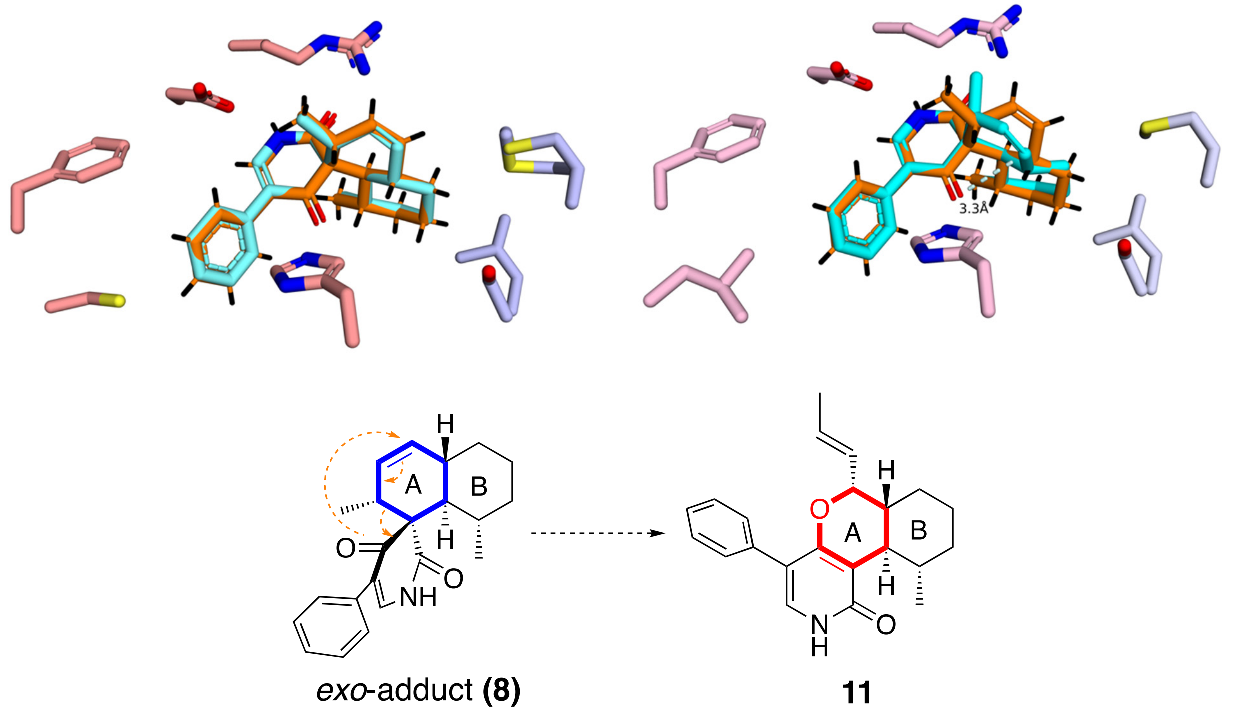


**Supplementary Figure 6 | Comparison of single crystal structure of 8 with alternative conformation of ligand 8 observed in the crystal structure of LepI-8 complex. (**a) Structure of **8** determined by single crystal X-ray analysis (colored in orange) is overlaid onto the conformer A of ligand **8** (colored in cyan). (b) Same structure is overlaid onto the conformer B of ligand **8**. (c) Theoretical retro-Claisen rearrangement of *exo*-adduct **8** would yield **11**. However, this reaction never occurred under our experimental condition. QM study suggests the barrier for this reaction (from **8** to transition state) is 10.7 kcal/mol higher compared to that from **9** to **TS-2.**

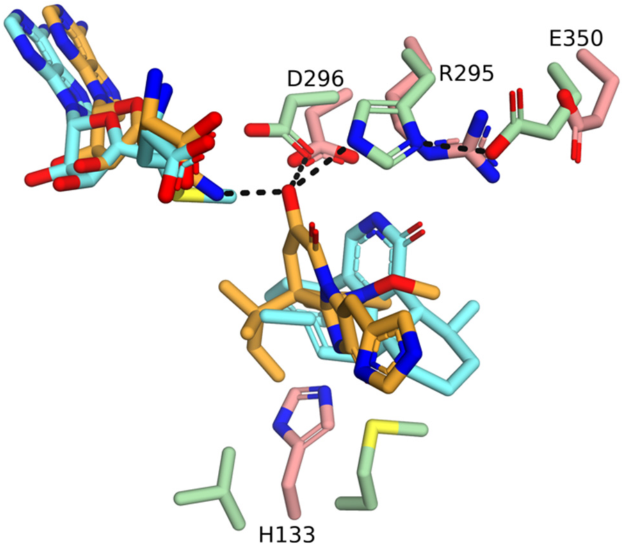


**Supplementary Figure 7 | Active site comparison of LepI with *O*-methyltransferase OxaC.** Crystal structure of OxaC pseudo Michalis-complex (ternary complex of OxaC-sinefungin-meleagrin, PDB entry 5W7S) is superimposed with LepI enzyme-product complex. Residues in OxaC and LepI are colored in pale green and salmon, respectively. Ligands in OxaC and LepI are colored in bright orange and cyan, respectively. The hydrogen bond networks in Oxac is shown in black dashes. The E350•R295 pair corresponds to a Glu-His dyad, and the role of His was established to be the general base which deprotonate the nucleophilic hydroxyl group for $S_N2$ attack at SAM. The D296 is conserved in both LepI and OxaC and its equivalent was proposed to position the substrate but not essential for catalysis. Remarkably, the H133 side is evolved substantially thus the main chain are not overlaid between LepI and OxaC. In OxaC, this side is taken by hydrophobic residues (Met and Val) which defined the hydrophobic pocket for substrate binding.

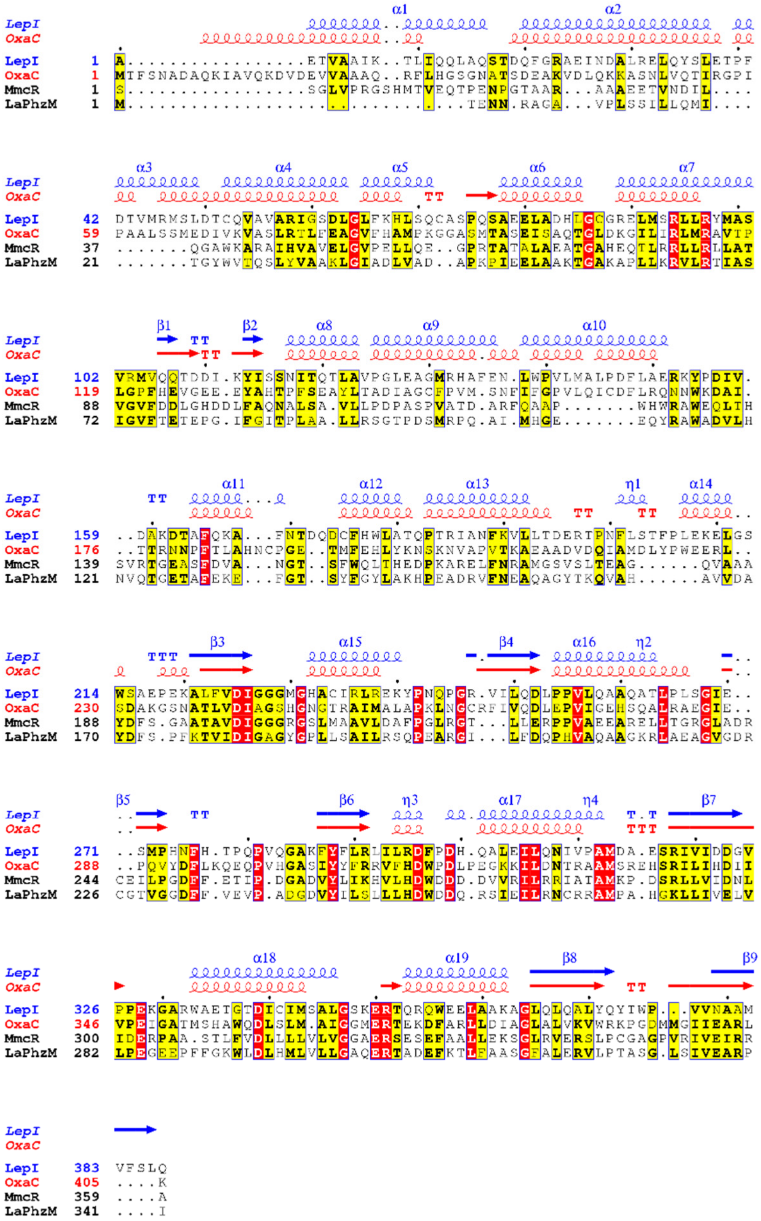

**Supplementary Figure 8 | Alignment of selected structural homologs of LepI identified by DALI search:** meleagrin O-methyltransferase OxaC (PDB: 5W7P); mitomycin 7-O-methyltransferase MmcR (PDB: 3GWZ); phenazine O-MT LaPhzM (PDB: 6C5B). The alignment was generated by T-coffee web-server using default parameters. The picture was generated by ESPript 3.0. Residues with strict identity are in white on a red background and those with identity above 70% are colored in red and framed in blue.

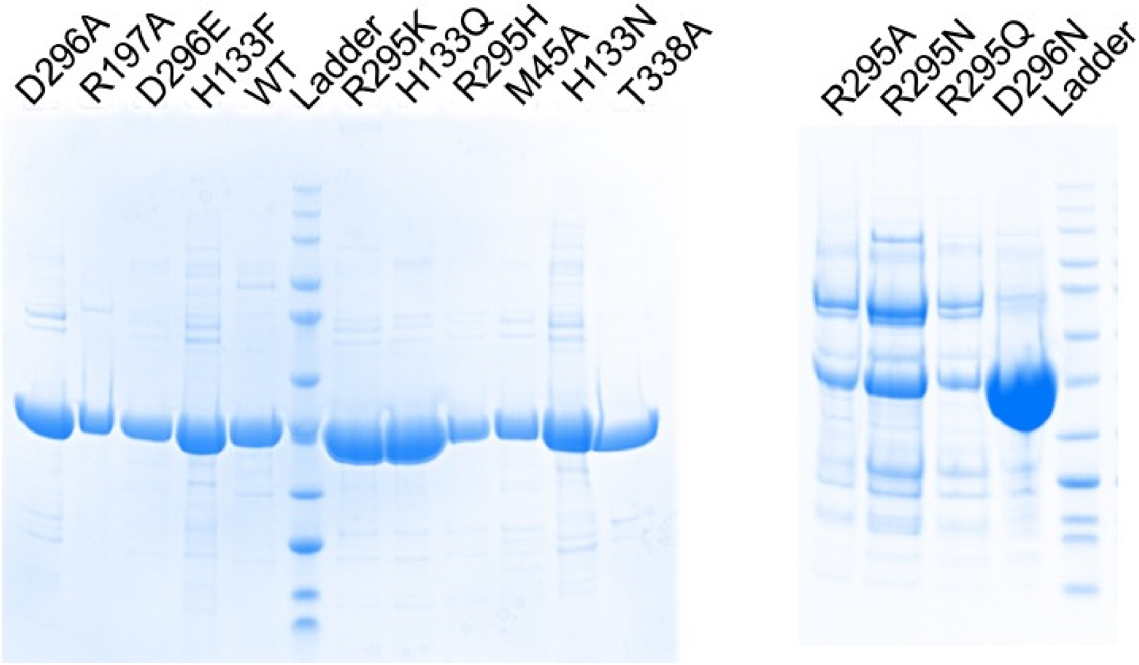


**Supplementary Figure 9 | SDS-PAGE analysis of purified LepI and mutant enzymes.**

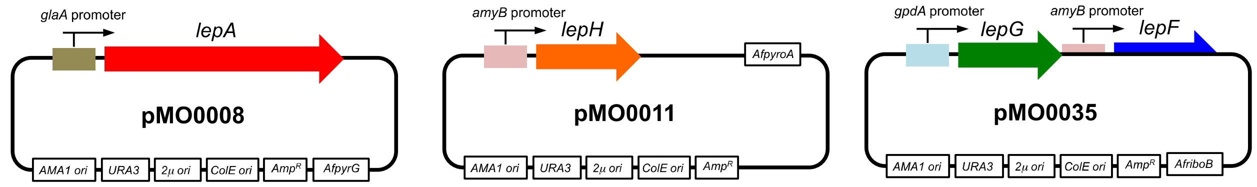


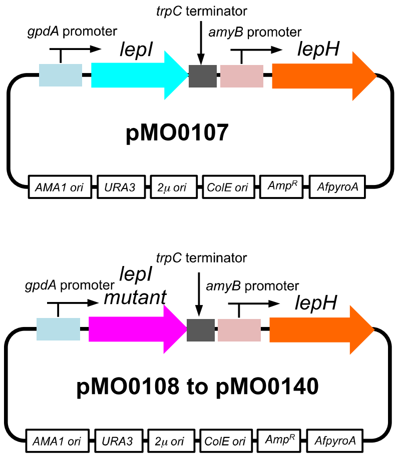


| Name | LepI mutant | Name | LepI mutant |
|---|---|---|---|
| pMO0107 | Wild type | pMO0122 | M45A |
| pMO0108 | D296N | pMO0125 | R197K |
| pMO0109 | D296A | pMO0126 | T338A |
| pMO0110 | D296E | pMO0127 | T338S |
| pMO0111 | R295K | pMO0133 | H133N |
| pMO0112 | R295E | pMO0134 | H133Q |
| pMO0113 | R295A | pMO0137 | R295Y |
| pMO0115 | R295H | pMO0138 | R295F |
| pMO0116 | H133A | pMO0139 | R295Q |
| pMO0117 | H133F | pMO0140 | R295N |
| pMO0121 | R197A | | |

**Supplementary Figure 10 | Plasmids used for heterologous expression of *lep* genes in *A. nidulans* A1145.**

**Supplementary Figure 11 |** $^{1}$H NMR spectrum of compound **8** in $CDCl_3$

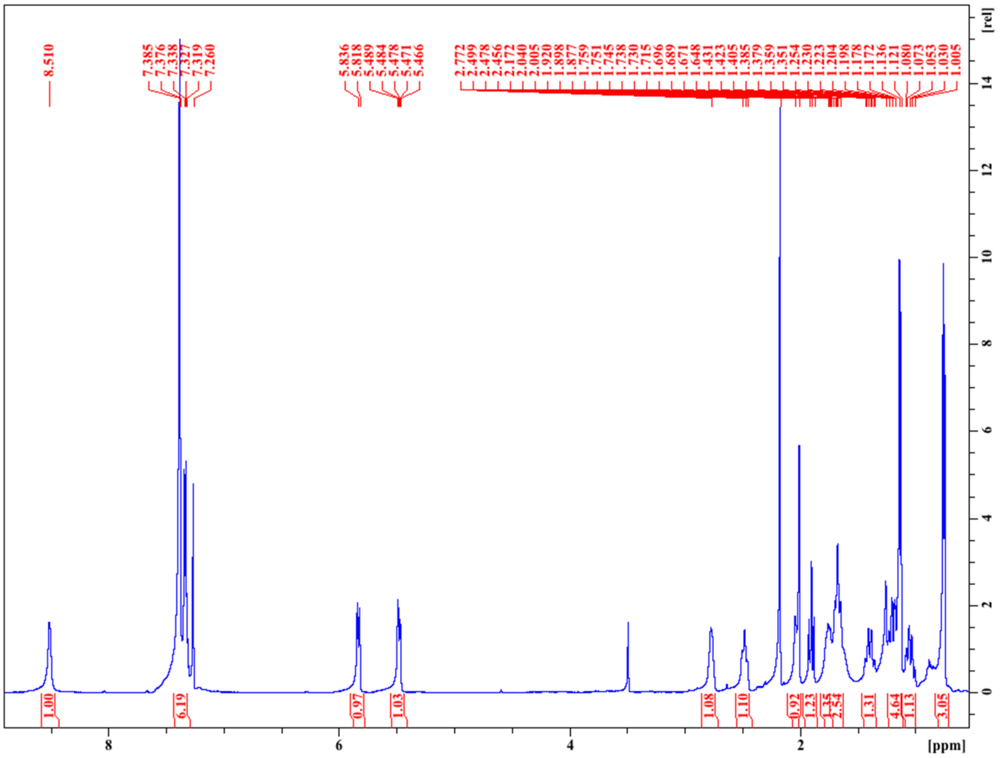

**Supplementary Figure 12 |** $^{13}$C NMR spectrum of compound **8** in $CDCl_3$

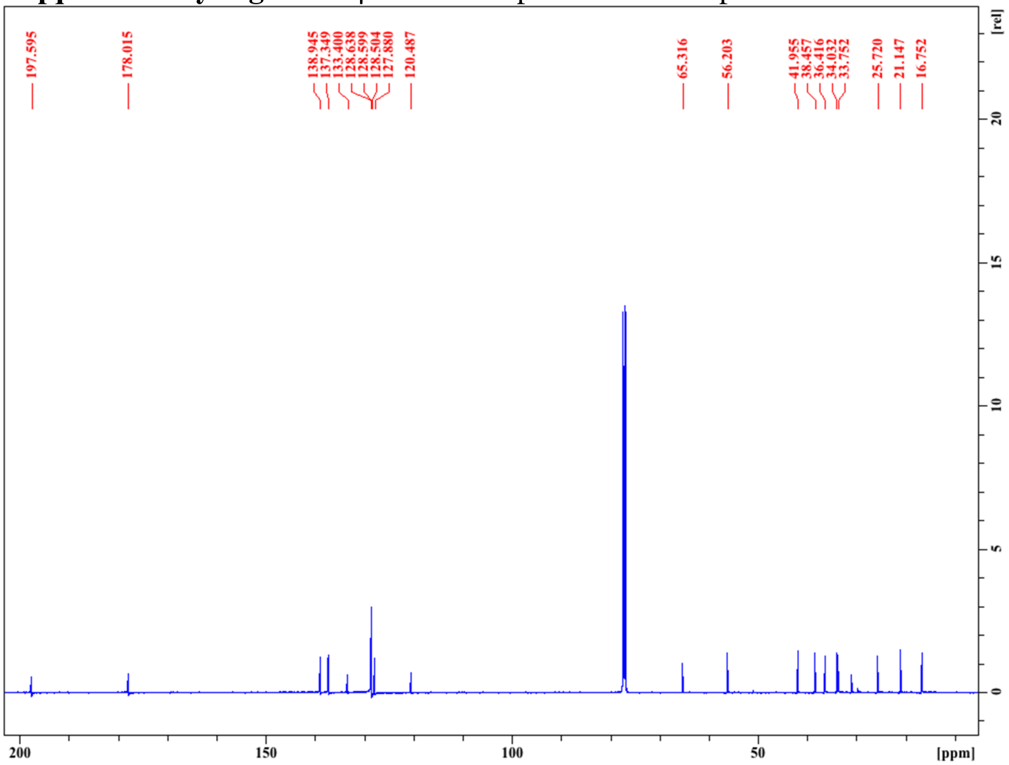

**Supplementary Figure 13 |** COSY spectrum of compound **8** in $CDCl_3$

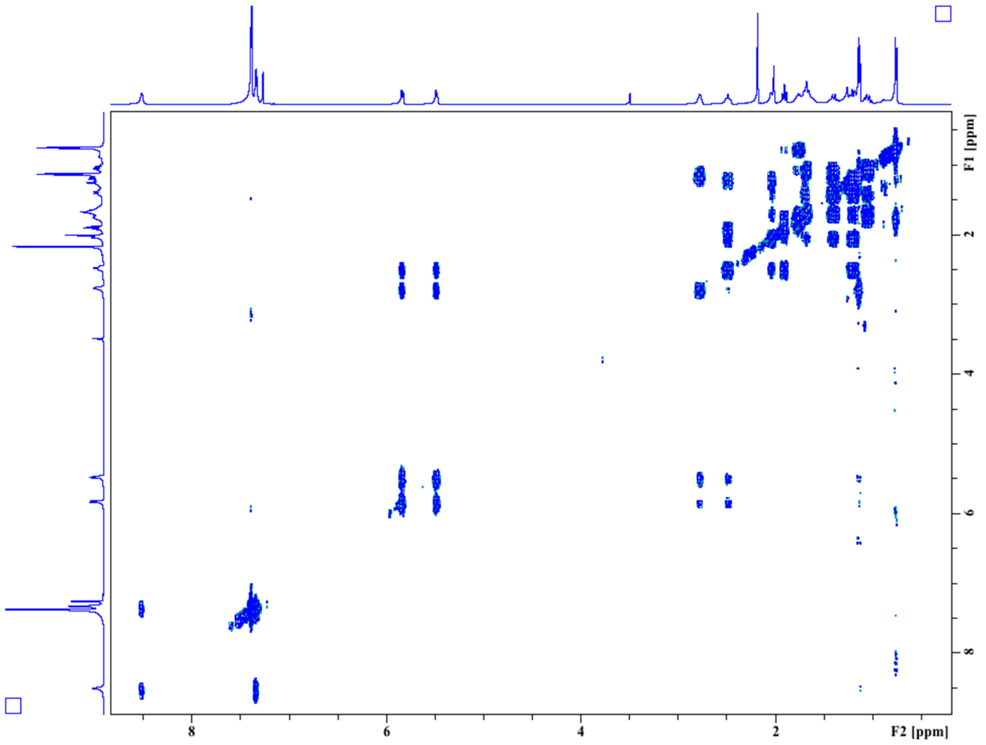


**Supplementary Figure 14 |** HSQC spectrum of compound **8** in $CDCl_3$

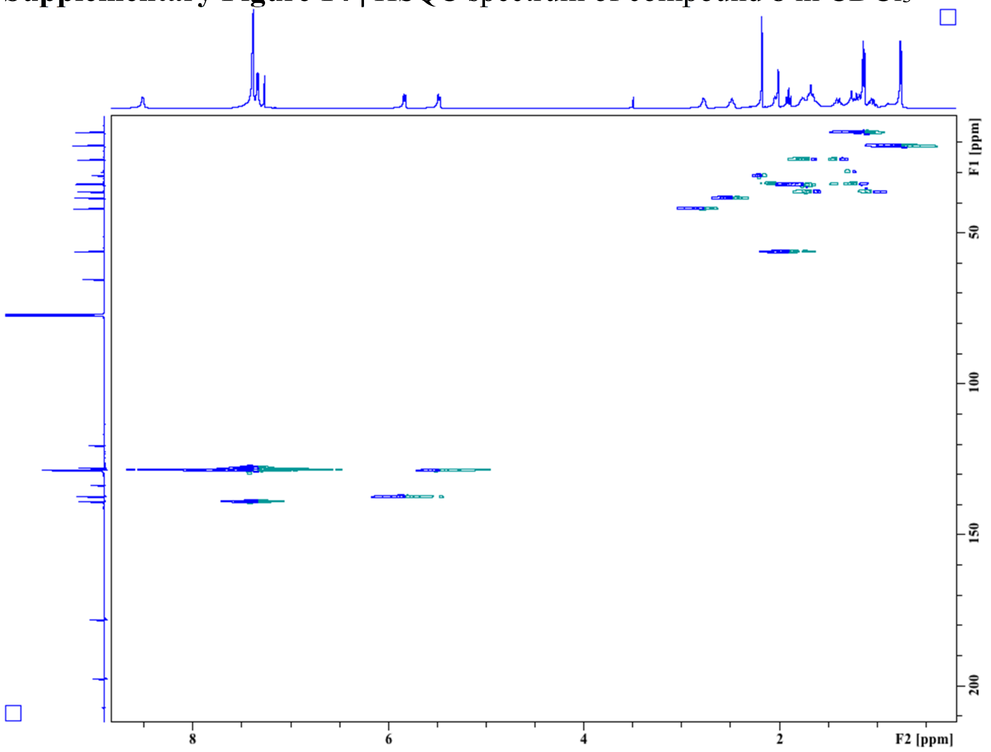

**Supplementary Figure 15 |** HMBC spectrum of compound **8** in $CDCl_3$

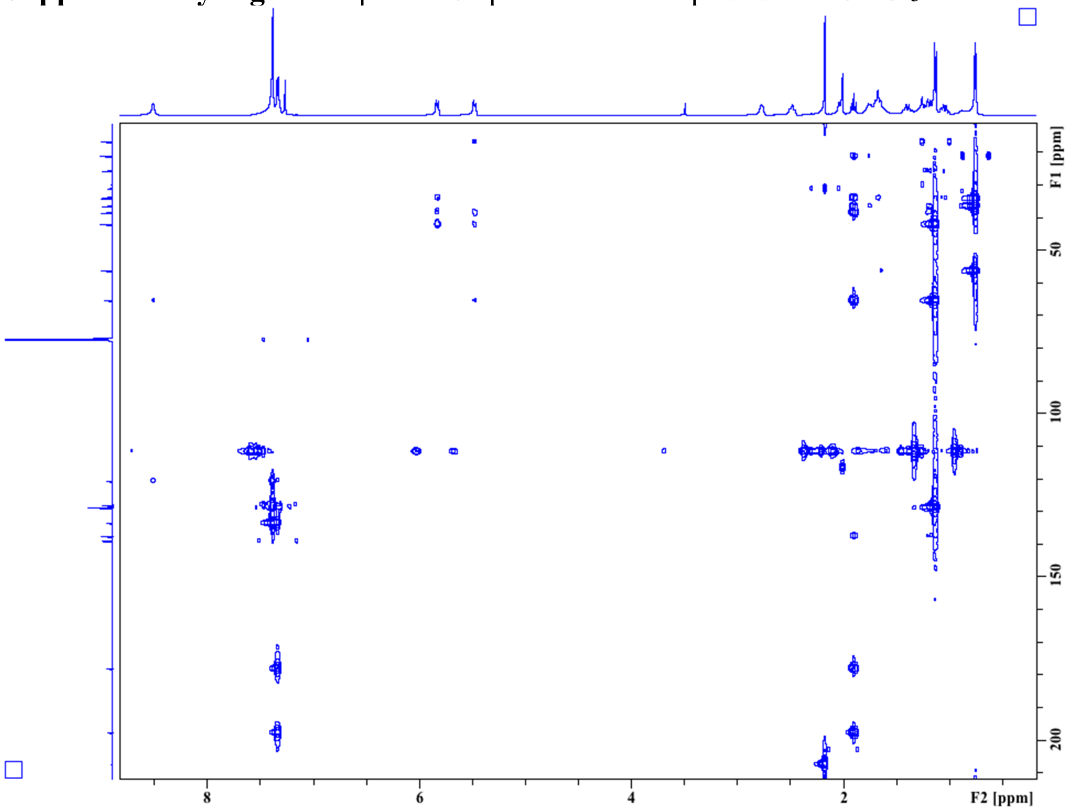

### SUPPLEMENTARY COMPUTATIONAL DATA

#### Cartesian coordinates of DFT optimized structures

**3**

E -1058.631935
G -1058.258012
0 1

| | | | |
|---|---|---|---|
| C | 0.086824 | -2.737225 | 0.864137 |
| C | 0.340197 | -1.709801 | 0.008793 |
| C | -0.792562 | -0.875269 | -0.443895 |
| C | -2.138153 | -1.173920 | 0.156473 |
| C | -2.314992 | -2.322180 | 1.100057 |
| C | -3.264315 | -0.456424 | -0.085402 |
| O | -0.642349 | 0.024755 | -1.272956 |
| O | -3.371475 | -2.664350 | 1.613154 |
| N | -1.151709 | -3.030582 | 1.379917 |
| C | 1.728736 | -1.466839 | -0.453217 |
| C | 2.015235 | -1.094016 | -1.778670 |
| C | 2.807040 | -1.652753 | 0.430862 |
| C | 3.335166 | -0.944078 | -2.203859 |
| C | 4.125599 | -1.519649 | -0.000805 |
| C | 4.396566 | -1.165200 | -1.323326 |
| C | -3.498391 | 0.760254 | -0.921472 |
| C | -3.666656 | 1.986581 | 0.023886 |
| C | -2.393073 | 2.385562 | 0.784563 |
| C | -1.317746 | 3.030873 | -0.115054 |
| C | -0.002071 | 3.196434 | 0.590760 |
| C | 1.101325 | 2.493191 | 0.283569 |
| C | 2.378094 | 2.596987 | 0.971707 |
| C | 3.466864 | 1.894302 | 0.617372 |
| C | 4.798450 | 1.950259 | 1.302473 |
| C | -4.770961 | 0.560531 | -1.767594 |
| H | 0.876072 | -3.410335 | 1.182122 |
| H | -4.146768 | -0.812883 | 0.450002 |
| H | -1.271486 | -3.821404 | 2.000099 |
| H | 1.200695 | -0.923402 | -2.470782 |
| H | 2.609296 | -1.880042 | 1.475065 |
| H | 3.533880 | -0.658482 | -3.233470 |
| H | 4.939691 | -1.670165 | 0.703224 |
| H | 5.422875 | -1.048097 | -1.660352 |
| H | -4.009747 | 2.831318 | -0.589898 |
| H | -4.477028 | 1.772174 | 0.735188 |
| H | -2.658168 | 3.083781 | 1.589871 |
| H | -1.964340 | 1.502691 | 1.274840 |
| H | -1.694261 | 4.006070 | -0.461328 |
| H | -1.164693 | 2.410206 | -1.002448 |
| H | 0.027580 | 3.905019 | 1.421825 |
| H | 1.049873 | 1.773817 | -0.534324 |
| H | 2.432785 | 3.283216 | 1.819689 |
| H | 3.393421 | 1.216433 | -0.231697 |
| H | 5.090504 | 0.956966 | 1.672028 |
| H | 4.793186 | 2.642704 | 2.151825 |
| H | 5.591497 | 2.267154 | 0.610162 |
| H | -4.676964 | -0.302893 | -2.435151 |
| H | -4.952750 | 1.447783 | -2.384162 |
| H | -5.651603 | 0.404205 | -1.131612 |
| H | -2.648054 | 0.929278 | -1.579866 |

**4**

E -1058.630796
G -1058.256608
0 1

| | | | |
|---|---|---|---|
| C | 1.522577 | -0.801476 | -1.648624 |
| C | 1.708967 | -0.717228 | -0.306623 |
| C | 0.526472 | -0.802084 | 0.574211 |
| C | -0.808231 | -1.014901 | -0.094704 |
| C | -0.917961 | -1.071332 | -1.570220 |
| C | -1.897330 | -1.092881 | 0.707619 |
| O | 0.609672 | -0.705031 | 1.797383 |
| O | -1.953741 | -1.196188 | -2.218321 |
| N | 0.294771 | -0.971712 | -2.246291 |
| C | 3.066996 | -0.513938 | 0.246941 |
| C | 4.170444 | -1.190568 | -0.299865 |
| C | 3.292826 | 0.390957 | 1.299720 |
| C | 5.462249 | -0.954812 | 0.169319 |
| C | 4.584448 | 0.621661 | 1.769864 |
| C | 5.674823 | -0.045167 | 1.206274 |
| C | -3.339487 | -1.269185 | 0.342839 |
| C | -4.268291 | -0.658035 | 1.410541 |
| C | -4.040797 | 0.824492 | 1.749174 |
| C | -4.178660 | 1.819970 | 0.569933 |
| C | -3.006934 | 1.827369 | -0.385248 |
| C | -1.793513 | 2.314527 | -0.073151 |
| C | -0.637424 | 2.267934 | -0.951929 |
| C | 1.805251 | 2.625664 | -1.492703 |
| C | -3.628522 | -2.778197 | 0.158975 |
| H | 2.352389 | -0.706709 | -2.340970 |
| H | -1.669463 | -1.041078 | 1.773110 |
| H | 0.219590 | -1.000449 | -3.255043 |
| H | 4.009615 | -1.925508 | -1.084886 |
| H | 2.450059 | 0.904396 | 1.745955 |
| H | 6.300114 | -1.492565 | -0.266820 |
| H | 4.739581 | 1.327735 | 2.581518 |
| H | 6.679689 | 0.135709 | 1.578403 |
| H | -3.520806 | -0.797225 | -0.623563 |
| H | -5.304282 | -0.793304 | 1.069918 |
| H | -4.171542 | -1.242445 | 2.337803 |
| H | -4.767234 | 1.104251 | 2.523864 |
| H | -3.047602 | 0.957522 | 2.198209 |
| H | -4.295091 | 2.824722 | 0.998372 |

| | | | |
|---|---|---|---|
| H | -5.107458 | 1.608603 | 0.024619 |
| H | -3.145804 | 1.370187 | -1.364620 |
| H | -1.634751 | 2.753896 | 0.914081 |
| H | -0.803089 | 1.867358 | -1.953500 |
| H | 0.757167 | 3.055261 | 0.403133 |
| H | 2.209299 | 3.634359 | -1.657937 |
| H | 1.568698 | 2.192539 | -2.471095 |
| H | 2.609360 | 2.032514 | -1.038212 |
| H | -3.401935 | -3.343232 | 1.071006 |
| H | -3.032676 | -3.180410 | -0.664618 |
| H | -4.688683 | -2.925449 | -0.077297 |
| C | 0.599431 | 2.665740 | -0.603991 |

***endo*-TS-1**

E -1058.612272

G -1058.238289

0 1

| | | | |
|---|---|---|---|
| C | -2.112688 | -0.520463 | -1.936037 |
| C | -0.865513 | -1.148621 | -2.143104 |
| C | -0.367865 | -2.148661 | -1.351045 |
| C | -0.787903 | -0.272635 | 0.848329 |
| C | -2.098767 | 0.247287 | 0.628258 |
| C | -2.941192 | -0.748580 | -0.824510 |
| C | -2.369297 | 1.729473 | 0.356813 |
| C | -3.874471 | 2.007692 | 0.192948 |
| C | -4.384632 | -0.255150 | -0.835229 |
| C | -4.523046 | 1.265549 | -0.979181 |
| C | -1.807714 | 2.566642 | 1.521310 |
| C | 0.993804 | -2.740111 | -1.509876 |
| C | -0.712699 | -1.480034 | 1.662448 |
| C | 1.717669 | -0.196987 | 0.718913 |
| C | 1.724611 | -1.239853 | 1.592290 |
| N | 0.579919 | -1.861734 | 2.030474 |
| O | -1.677552 | -2.162804 | 2.025703 |
| C | 0.410633 | 0.346029 | 0.275686 |
| O | 0.331720 | 1.278301 | -0.543417 |
| C | 2.989752 | 0.355935 | 0.202076 |
| C | 4.067878 | -0.496656 | -0.095473 |
| C | 3.164176 | 1.737773 | 0.005351 |
| C | 5.287605 | 0.010354 | -0.542674 |
| C | 4.381456 | 2.242329 | -0.447925 |
| C | 5.450472 | 1.384954 | -0.719770 |
| H | -2.798377 | -0.174253 | 1.354526 |
| H | -2.817326 | -1.705255 | -0.318502 |
| H | -1.015443 | -2.634206 | -0.626735 |
| H | -1.842447 | 2.023200 | -0.552652 |
| H | -4.394505 | 1.735098 | 1.124804 |
| H | -4.016663 | 3.088500 | 0.065167 |
| H | -4.925598 | -0.766091 | -1.643119 |
| H | -4.863375 | -0.567800 | 0.101507 |
| H | -5.587149 | 1.524981 | -1.043295 |
| H | -4.069022 | 1.600163 | -1.922200 |
| H | -2.243301 | 2.254868 | 2.479145 |
| H | -0.721858 | 2.467268 | 1.583996 |
| H | -2.045753 | 3.626871 | 1.372994 |
| H | 1.491966 | -2.820581 | -0.536842 |
| H | 0.923681 | -3.761406 | -1.912789 |
| H | 1.628631 | -2.143918 | -2.172107 |
| H | -0.225701 | -0.744679 | -2.924858 |
| H | -2.379438 | 0.279910 | -2.622203 |
| H | 2.648959 | -1.649938 | 1.986129 |
| H | 0.631795 | -2.664739 | 2.643472 |
| H | 3.940317 | -1.571995 | 0.003030 |
| H | 2.336585 | 2.407712 | 0.201765 |
| H | 6.104704 | -0.670946 | -0.766476 |
| H | 4.495966 | 3.314196 | -0.588541 |
| H | 6.396966 | 1.783600 | -1.075413 |

**TS-2**

E -1058.61584

G -1058.236209

0 1

| | | | |
|---|---|---|---|
| C | 0.854209 | 0.664410 | -0.627792 |
| C | 0.316138 | 1.794536 | 1.811465 |
| C | 0.489584 | 0.550716 | 2.402521 |
| C | 1.565563 | -0.269049 | 2.079126 |
| C | 2.645852 | 0.179998 | 1.186205 |
| C | 2.221680 | 0.122162 | -0.387318 |
| C | 4.004504 | -0.511100 | 1.421843 |
| C | 4.100751 | -1.909598 | 0.808604 |
| C | 2.447410 | -1.286789 | -1.013928 |
| C | 3.847477 | -1.834705 | -0.699515 |
| C | 2.219588 | -1.226066 | -2.531984 |
| C | 0.724516 | 1.947364 | -1.285483 |
| C | -0.293037 | -0.146173 | -0.299882 |
| C | -1.696340 | 1.526854 | -1.362025 |
| O | 1.647915 | 2.734089 | -1.546169 |
| O | -0.132648 | -1.209638 | 0.359066 |
| N | -0.588439 | 2.290135 | -1.635787 |
| C | -1.629834 | 0.352667 | -0.673221 |
| C | -2.861949 | -0.382923 | -0.310293 |
| C | -2.927972 | -1.786918 | -0.383685 |
| C | -4.009598 | 0.314112 | 0.111859 |
| C | -4.108163 | -2.461522 | -0.068757 |
| C | -5.192236 | -0.361415 | 0.421394 |
| C | -5.247219 | -1.754747 | 0.331209 |
| C | -0.949575 | 2.578385 | 1.895387 |
| H | 2.795362 | 1.252838 | 1.344755 |
| H | 2.932734 | 0.813689 | -0.854903 |
| H | 4.780902 | 0.121322 | 0.969251 |
| H | 4.215010 | -0.536421 | 2.498895 |
| H | 3.362317 | -2.581134 | 1.271587 |
| H | 5.089961 | -2.336872 | 1.017691 |
| H | 1.717178 | -1.974020 | -0.579828 |
| H | 3.949037 | -2.829802 | -1.153360 |
| H | 4.614036 | -1.196186 | -1.167975 |

| | | | |
|---|---|---|---|
| H | 1.188537 | -0.945294 | -2.772763 |
| H | 2.885860 | -0.490227 | -3.002360 |
| H | -0.335817 | 0.121284 | 2.965743 |
| H | 1.599116 | -1.274368 | 2.489885 |
| H | -2.636340 | 1.929962 | -1.725564 |
| H | -2.047586 | -2.343113 | -0.683910 |
| H | -3.969581 | 1.395958 | 0.218397 |
| H | -4.138182 | -3.546360 | -0.136982 |
| H | -6.064798 | 0.200683 | 0.745872 |
| H | -6.163967 | -2.284651 | 0.577410 |
| H | -0.678913 | 3.153411 | -2.158375 |
| H | -1.791178 | 1.972657 | 2.243873 |
| H | -1.199881 | 2.999362 | 0.914214 |
| H | -0.818996 | 3.433641 | 2.576141 |
| H | 1.165054 | 2.318996 | 1.384922 |
| H | 2.418978 | -2.203100 | -2.989666 |

***exo*-TS-3**

E -1058.612601

G -1058.238776

0 1

| | | | |
|---|---|---|---|
| C | -2.824871 | 1.808363 | -0.730353 |
| C | -1.656919 | 2.569138 | -0.956026 |
| C | -0.480068 | 2.051921 | -1.424520 |
| C | -0.737211 | 0.106036 | 0.735841 |
| C | -2.025308 | -0.476972 | 0.537410 |
| C | -2.948829 | 0.427656 | -0.960530 |
| C | -2.185013 | -1.964467 | 0.200779 |
| C | -3.661687 | -2.368889 | 0.062294 |
| C | -4.355171 | -0.158778 | -0.976645 |
| C | -4.376611 | -1.683068 | -1.102137 |
| C | -1.519907 | -2.798313 | 1.314525 |
| C | 0.784447 | 2.835053 | -1.549328 |
| C | -0.730632 | 1.278967 | 1.593881 |
| C | 1.763372 | 0.179111 | 0.620384 |
| C | 1.707382 | 1.212116 | 1.507722 |
| N | 0.532067 | 1.767072 | 1.940983 |
| O | -1.745431 | 1.855115 | 2.005131 |
| C | 0.493709 | -0.401811 | 0.122866 |
| O | 0.476192 | -1.264855 | -0.773498 |
| C | 3.070566 | -0.310867 | 0.124716 |
| C | 4.131126 | 0.583182 | -0.106621 |
| C | 3.295329 | -1.678852 | -0.112689 |
| C | 5.380070 | 0.129191 | -0.529512 |
| C | 4.541924 | -2.130729 | -0.540726 |
| C | 5.591982 | -1.232700 | -0.747243 |
| H | -2.727401 | -0.142226 | 1.301298 |
| H | -2.282673 | 0.003893 | -1.713067 |
| H | -0.445960 | 1.036427 | -1.807980 |
| H | -1.658585 | -2.174826 | -0.733227 |
| H | -4.189490 | -2.141914 | 1.001698 |
| H | -3.714704 | -3.457577 | -0.066573 |
| H | -4.905964 | 0.283931 | -1.818167 |
| H | -4.883825 | 0.148701 | -0.063342 |
| H | -5.416241 | -2.028628 | -1.162413 |
| H | -3.894874 | -1.974133 | -2.046995 |
| H | -1.954038 | -2.564625 | 2.294716 |
| H | -0.444744 | -2.612294 | 1.356816 |
| H | -1.673540 | -3.867277 | 1.124205 |
| H | 1.082885 | 2.917838 | -2.604007 |
| H | 1.605626 | 2.317114 | -1.038537 |
| H | 0.688608 | 3.843002 | -1.132970 |
| H | -1.670612 | 3.607661 | -0.630388 |
| H | -3.660991 | 2.317565 | -0.255716 |
| H | 2.606458 | 1.661089 | 1.917304 |
| H | 0.532932 | 2.553046 | 2.577223 |
| H | 3.970049 | 1.650537 | 0.025994 |
| H | 2.482855 | -2.380272 | 0.030520 |
| H | 6.182000 | 0.842520 | -0.702508 |
| H | 4.694062 | -3.193048 | -0.714080 |
| H | 6.561541 | -1.589818 | -1.084289 |

**TS-4**

E -1058.600898

G -1058.219935

0 1

| | | | |
|---|---|---|---|
| O | -0.540037 | -1.325593 | -0.198057 |
| O | 1.700758 | 2.834212 | -0.123504 |
| N | -0.565695 | 2.722053 | -0.317772 |
| H | -0.543505 | 3.733005 | -0.356276 |
| C | 0.648108 | 0.575923 | 2.472409 |
| H | -0.169917 | 1.214442 | 2.809267 |
| C | 0.344960 | -0.779136 | 2.368429 |
| H | -0.610253 | -1.095929 | 2.785149 |
| C | 0.935947 | -1.690462 | 1.504867 |
| H | 0.579024 | -2.714761 | 1.566257 |
| C | 1.988014 | -1.459913 | 0.465303 |
| H | 1.705358 | -2.183367 | -0.306744 |
| C | 3.442101 | -1.836388 | 0.839521 |
| H | 3.885009 | -1.077291 | 1.496904 |
| H | 3.466311 | -2.786616 | 1.387251 |
| C | 4.249669 | -1.962481 | -0.464638 |
| H | 3.887806 | -2.848407 | -1.005582 |
| H | 5.307738 | -2.145742 | -0.238613 |
| C | 4.116388 | -0.729008 | -1.375080 |
| H | 4.607846 | -0.932119 | -2.335606 |
| H | 4.660022 | 0.114417 | -0.923619 |
| C | 2.662106 | -0.269610 | -1.625188 |
| H | 2.116167 | -1.070084 | -2.145847 |
| C | 1.993703 | -0.053725 | -0.228817 |
| H | 2.690038 | 0.607491 | 0.294051 |
| C | 0.670694 | 0.669744 | -0.224623 |
| C | -0.557408 | -0.057853 | -0.292389 |
| C | -1.824364 | 0.687175 | -0.326425 |
| C | -1.755609 | 2.048076 | -0.375513 |

| | | | |
|---|---|---|---|
| H | -2.640748 | 2.668670 | -0.466198 |
| C | 0.692870 | 2.115793 | -0.222621 |
| C | -3.132983 | -0.003694 | -0.316474 |
| C | -3.338501 | -1.200908 | -1.025645 |
| H | -2.510905 | -1.643828 | -1.565982 |
| C | -4.588060 | -1.817786 | -1.032534 |
| H | -4.725176 | -2.740104 | -1.591425 |
| C | -5.660675 | -1.259518 | -0.333026 |
| H | -6.632944 | -1.745166 | -0.339897 |
| C | -5.468672 | -0.077958 | 0.384456 |
| H | -6.289453 | 0.359762 | 0.947163 |
| C | -4.216863 | 0.536321 | 0.398459 |
| H | -4.068895 | 1.435603 | 0.991487 |
| C | 1.994840 | 1.204304 | 2.596392 |
| H | 2.077160 | 1.598892 | 3.620185 |
| H | 2.126777 | 2.056880 | 1.915644 |
| H | 2.805409 | 0.490023 | 2.452131 |
| C | 2.643966 | 0.977893 | -2.514273 |
| H | 3.096003 | 1.832823 | -2.003220 |
| H | 1.619566 | 1.258129 | -2.781011 |
| H | 3.194288 | 0.789228 | -3.444120 |

***endo*-TS-5 (Ambimodal (Z)-QM TS to 6 and 7)**

E -1058.614067

G -1058.237248

0 1

| | | | |
|---|---|---|---|
| C | -0.780652 | 0.700287 | -0.158592 |
| C | -0.295193 | -0.751124 | 2.369077 |
| C | -0.270084 | -1.863914 | 1.577604 |
| C | -1.277052 | -2.203008 | 0.643681 |
| C | -2.454329 | -1.466687 | 0.425616 |
| C | -1.957554 | 0.096395 | -0.689848 |
| C | -3.582974 | -2.144077 | -0.340869 |
| C | -4.768294 | -1.214948 | -0.609640 |
| C | -3.273685 | 0.865070 | -0.836916 |
| C | -4.377605 | -0.006309 | -1.460594 |
| C | -3.033239 | 2.104607 | -1.722841 |
| C | -0.817354 | 1.840167 | 0.743580 |
| C | 0.481537 | 0.063385 | -0.536464 |
| C | 1.641039 | 1.851125 | 0.631133 |
| O | -1.817916 | 2.342906 | 1.266943 |
| O | 0.462052 | -1.011466 | -1.170668 |
| N | 0.440226 | 2.392265 | 1.021467 |
| C | 1.736791 | 0.715455 | -0.111675 |
| C | 3.061911 | 0.163370 | -0.472862 |
| C | 3.300537 | -1.222920 | -0.463698 |
| C | 4.129940 | 1.017345 | -0.799833 |
| C | 4.568008 | -1.727667 | -0.748892 |
| C | 5.398807 | 0.510853 | -1.078239 |
| C | 5.624820 | -0.866231 | -1.051860 |
| C | 0.834879 | -0.325160 | 3.246530 |
| H | -2.795974 | -0.855242 | 1.261637 |
| H | -1.713751 | -0.507741 | -1.563974 |
| H | -3.190272 | -2.537054 | -1.289161 |
| H | -3.924892 | -3.013112 | 0.238092 |
| H | -5.178983 | -0.868486 | 0.350113 |
| H | -5.568451 | -1.778912 | -1.105154 |
| H | -3.595683 | 1.215377 | 0.147820 |
| H | -5.261748 | 0.620791 | -1.632532 |
| H | -4.044062 | -0.350814 | -2.451693 |
| H | -2.341800 | 2.801718 | -1.245717 |
| H | -2.625003 | 1.816118 | -2.699371 |
| H | 0.633620 | -2.470619 | 1.573195 |
| H | -1.085917 | -3.056642 | -0.001670 |
| H | 2.519752 | 2.375355 | 0.992336 |
| H | 2.481386 | -1.896568 | -0.246270 |
| H | 3.956542 | 2.088918 | -0.860974 |
| H | 4.730924 | -2.802490 | -0.733935 |
| H | 6.206730 | 1.192829 | -1.331135 |
| H | 6.610887 | -1.264459 | -1.276057 |
| H | 0.417369 | 3.192665 | 1.640251 |
| H | 0.537987 | -0.355688 | 4.304566 |
| H | 1.113171 | 0.714345 | 3.031799 |
| H | 1.721118 | -0.952345 | 3.110449 |
| H | -1.199152 | -0.151805 | 2.435677 |
| H | -3.979993 | 2.630418 | -1.895138 |

***exo*-TS-6**

**5**

E -1058.659236

G -1058.275957

0 1

| | | | |
|---|---|---|---|
| C | 3.855701 | 1.050233 | -0.246211 |
| C | 3.340252 | 2.481114 | -0.428392 |
| C | 1.823167 | 2.473601 | -0.651094 |
| C | 1.054482 | 1.823359 | 0.518979 |
| C | 1.566485 | 0.391933 | 0.863575 |
| C | 3.116947 | 0.307243 | 0.894062 |
| C | 0.841530 | -0.770477 | 0.068390 |
| C | 1.346541 | -2.149464 | 0.622350 |
| C | 3.608204 | -1.109363 | 1.003999 |
| C | 2.834436 | -2.184184 | 0.858717 |
| C | 1.079636 | 2.708890 | 1.778326 |
| C | 0.876081 | -3.336060 | -0.236642 |
| C | -0.662138 | -0.648462 | 0.464737 |
| C | 1.117434 | -0.700537 | -1.431120 |
| C | -1.241708 | -0.130930 | -1.834356 |
| O | -0.968142 | -0.838589 | 1.636441 |
| O | 2.184236 | -0.978010 | -1.952658 |
| C | -1.658733 | -0.266251 | -0.549374 |
| N | 0.058966 | -0.306740 | -2.244882 |
| C | -3.071456 | -0.006569 | -0.184683 |
| C | -3.759152 | -0.830604 | 0.722812 |

C -3.762084 1.071819 -0.763784
C -5.099285 -0.590419 1.020623
C -5.104784 1.306041 -0.469624
C -5.779834 0.473599 0.424205
H 1.608253 1.930626 -1.580852
H 3.842390 2.956436 -1.280480
H 3.586637 3.087074 0.454409
H 4.929109 1.056988 -0.014049
H 3.736953 0.493295 -1.178485
H 0.002234 1.753136 0.216020
H 1.236795 0.179208 1.886781
H 3.405250 0.823859 1.824469
H 1.449009 3.495496 -0.797190
H 0.852019 -2.240853 1.598136
H 2.082110 2.781487 2.213859
H 0.412948 2.300829 2.546216
H 0.745210 3.726804 1.545412
H 1.366836 -3.340275 -1.214660
H 1.116627 -4.280499 0.263873
H -0.210063 -3.309249 -0.388417
H -1.936914 0.117851 -2.629742
H 0.280470 -0.287715 -3.233143
H -3.237222 -1.653833 1.194716
H -3.233196 1.748343 -1.430865
H -5.615258 -1.240585 1.722213
H -5.617488 2.148068 -0.927241
H -6.824122 0.657855 0.661650
H 3.273472 -3.178452 0.936685
H 4.676095 -1.232997 1.181620

**6**
E -1058.661359
G -1058.277534
0 1
C 0.706580 -0.544975 -0.121346
C 1.938347 0.332586 0.239535
C 3.266561 -0.461336 0.334236
C 3.118492 -1.830698 0.923114
C 1.953707 -2.457269 1.075598
C 0.603526 -1.840060 0.818473
C 4.320106 0.399211 1.057408
C 2.200703 1.550692 -0.698060
C 3.272899 2.442439 -0.034977
C 4.576592 1.691516 0.270804
C 0.986800 2.402234 -1.091431
C -0.084938 -1.590631 2.180827
C -0.614530 0.213072 0.166132
C 0.809998 -1.078654 -1.548683
C -1.816592 -0.217625 -0.578022
C -1.634787 -0.952262 -1.705534
O -0.657279 1.049890 1.057288
O 1.851056 -1.313021 -2.137504
N -0.404112 -1.362338 -2.169576

C -3.180198 0.143620 -0.126117
C -3.484115 1.427104 0.360772
C -4.211918 -0.809612 -0.189407
C -4.784157 1.743795 0.751675
C -5.512531 -0.487239 0.195416
C -5.804121 0.793381 0.667651
H 1.710096 0.725849 1.236999
H -0.018326 -2.568017 0.278400
H 3.631603 -0.617548 -0.690615
H 5.254263 -0.167485 1.169519
H 5.269616 2.344115 0.816950
H 3.967880 0.633821 2.072284
H 5.069076 1.432627 -0.677681
H 2.623521 1.146237 -1.628298
H 2.854592 2.850819 0.897313
H 3.485093 3.303120 -0.683044
H 4.043043 -2.349809 1.176398
H 1.927467 -3.470002 1.476908
H 1.307890 3.207614 -1.762860
H 0.502779 2.850115 -0.219128
H 0.227112 1.822686 -1.630363
H 0.009805 -2.493939 2.793654
H -1.149412 -1.361803 2.079222
H 0.390962 -0.764803 2.716061
H -2.697897 2.167513 0.433403
H -3.984074 -1.820092 -0.520208
H -5.001388 2.742301 1.121875
H -6.293171 -1.241505 0.140500
H -6.815125 1.045969 0.975816
H -0.322111 -1.754464 -3.099821
H -2.473247 -1.237647 -2.332882

**7**
E -1058.658332
G -1058.276639
0 1
C 0.467372 1.710001 -0.519054
C 1.822125 1.286255 0.093483
C 2.212848 -0.103583 -0.485083
C 3.673500 -0.528093 -0.197205
C 2.877619 2.375208 -0.149640
C 4.261043 1.913545 0.312245
C 4.652259 0.637529 -0.435920
C 4.085984 -1.721367 -1.071220
C -0.303633 2.646716 0.375965
C -1.574664 2.502658 0.753493
C -2.305572 3.443131 1.667530
O -0.372734 0.585423 -0.901855
C -0.178333 -0.609263 -0.287419
C 1.106159 -1.049485 -0.031445
C 1.295439 -2.261322 0.745475
C -1.367005 -1.356883 0.039023
C -1.159311 -2.542191 0.688471

| | | | |
|---|---|---|---|
| N | 0.094028 | -2.954185 | 1.013279 |
| O | 2.343775 | -2.745374 | 1.179410 |
| C | -2.738142 | -0.879580 | -0.254295 |
| C | -3.742480 | -0.979886 | 0.723470 |
| C | -3.074684 | -0.326661 | -1.502074 |
| C | -5.045068 | -0.555985 | 0.460524 |
| C | -4.374337 | 0.104131 | -1.760678 |
| C | -5.366032 | -0.010290 | -0.783164 |
| H | 1.699237 | 1.165491 | 1.178315 |
| H | 0.673809 | 2.202992 | -1.478748 |
| H | 2.125037 | -0.009367 | -1.582307 |
| H | 5.659776 | 0.313353 | -0.145450 |
| H | 3.744386 | -0.827967 | 0.853200 |
| H | 4.697951 | 0.859408 | -1.513956 |
| H | 2.920405 | 2.606542 | -1.224729 |
| H | 2.584124 | 3.302113 | 0.359966 |
| H | 4.245847 | 1.721589 | 1.394806 |
| H | 5.001105 | 2.705127 | 0.140095 |
| H | 4.021989 | -1.458617 | -2.136247 |
| H | 5.123856 | -2.006791 | -0.861288 |
| H | 3.460335 | -2.594365 | -0.882803 |
| H | 0.273297 | 3.502868 | 0.728469 |
| H | -2.141393 | 1.652014 | 0.383450 |
| H | -3.193597 | 3.859647 | 1.173149 |
| H | -1.673286 | 4.276926 | 1.992723 |
| H | -2.665490 | 2.918724 | 2.563214 |
| H | -1.974317 | -3.199460 | 0.970466 |
| H | 0.235351 | -3.818549 | 1.522304 |
| H | -3.489699 | -1.369143 | 1.706329 |
| H | -2.309277 | -0.231044 | -2.263865 |
| H | -5.804422 | -0.638615 | 1.233735 |
| H | -4.614285 | 0.529053 | -2.731779 |
| H | -6.377784 | 0.329464 | -0.987328 |

**8-halfchair**

E -1058.660373

G -1058.276627

0 1

| | | | |
|---|---|---|---|
| C | 0.675910 | 1.493272 | 1.486691 |
| N | -0.552611 | 2.273143 | -1.344493 |
| O | -0.467274 | -1.333955 | 0.466016 |
| C | 1.197671 | 0.646987 | 2.618690 |
| O | 1.700699 | 2.408185 | -1.380575 |
| C | 1.993580 | -0.410322 | 2.475068 |
| C | 2.368471 | -1.006746 | 1.153960 |
| C | 3.844013 | -1.443787 | 1.079926 |
| C | 4.093752 | -2.245254 | -0.205000 |
| C | 3.605416 | -1.496187 | -1.453786 |
| C | 2.146777 | -1.003284 | -1.333357 |
| C | 2.047386 | -0.104349 | -0.063901 |
| C | 0.733094 | 0.718154 | 0.085791 |
| C | -0.524332 | -0.180819 | 0.061371 |
| C | -1.794433 | 0.428639 | -0.396621 |
| C | -1.725856 | 1.593435 | -1.089717 |
| C | 0.709424 | 1.838059 | -0.959261 |
| C | -3.104983 | -0.214700 | -0.141795 |
| C | -3.288522 | -1.602496 | -0.272840 |
| C | -4.542256 | -2.172663 | -0.057135 |
| C | -5.635620 | -1.376966 | 0.291442 |
| C | -5.464228 | 0.000959 | 0.432792 |
| C | -4.209932 | 0.573231 | 0.225925 |
| C | 1.423857 | 2.846340 | 1.510289 |
| H | -0.384532 | 1.712115 | 1.676291 |
| H | -0.551362 | 3.042983 | -2.002744 |
| H | 0.932382 | 0.999459 | 3.614900 |
| H | 2.354672 | -0.933435 | 3.360672 |
| H | 1.760692 | -1.916995 | 1.059288 |
| H | 4.099868 | -2.050453 | 1.959002 |
| H | 4.491941 | -0.555638 | 1.108815 |
| H | 3.555268 | -3.201146 | -0.131328 |
| H | 5.158529 | -2.491620 | -0.304829 |
| H | 3.701842 | -2.144769 | -2.334617 |
| H | 4.250877 | -0.624143 | -1.637540 |
| H | 1.505607 | -1.876557 | -1.152304 |
| H | 2.837883 | 0.652854 | -0.135140 |
| H | -2.613474 | 2.049728 | -1.515903 |
| H | -2.445677 | -2.227120 | -0.538977 |
| H | -4.665085 | -3.247065 | -0.166208 |
| H | -6.610098 | -1.827558 | 0.459640 |
| H | -6.302649 | 0.630398 | 0.719560 |
| H | -4.076255 | 1.641837 | 0.376401 |
| H | 2.463137 | 2.727724 | 1.193809 |
| H | 0.954314 | 3.596970 | 0.868114 |
| H | 1.421075 | 3.232858 | 2.535493 |
| C | 1.692167 | -0.380860 | -2.659023 |
| H | 2.273783 | 0.509630 | -2.913566 |
| H | 1.807836 | -1.112974 | -3.467162 |
| H | 0.632169 | -0.096789 | -2.640809 |

**8-boat**

E -1058.659973

G -1058.276856

0 1

| | | | |
|---|---|---|---|
| C | -1.044731 | 1.742044 | -1.217831 |
| O | 0.474960 | -0.801086 | -1.232758 |
| C | -2.504282 | 1.996516 | -1.476652 |
| O | -1.734736 | 2.184516 | 1.770739 |
| C | -3.347768 | 0.964472 | -1.493013 |
| C | -2.855759 | -0.405582 | -1.094682 |
| C | -4.033150 | -1.375717 | -0.904262 |
| C | -3.590260 | -2.722789 | -0.333867 |
| C | -2.861353 | -2.515329 | 0.995894 |
| C | -1.624062 | -1.611238 | 0.842163 |
| C | -2.021907 | -0.251019 | 0.214286 |

| | | | |
|---|---|---|---|
| C | -0.816621 | 0.756433 | 0.033060 |
| C | 0.495728 | 0.023378 | -0.323888 |
| C | 1.740359 | 0.386093 | 0.378467 |
| C | 1.663246 | 1.273532 | 1.405207 |
| C | -0.760395 | 1.643927 | 1.275653 |
| C | 3.045142 | -0.191053 | -0.021620 |
| C | 3.172800 | -1.552868 | -0.345065 |
| C | 4.416025 | -2.086767 | -0.679000 |
| C | 5.554037 | -1.277103 | -0.699782 |
| C | 5.438150 | 0.079024 | -0.391216 |
| C | 4.194154 | 0.616601 | -0.062784 |
| C | -0.930586 | -1.440849 | 2.202028 |
| H | -0.667881 | 1.155705 | -2.065005 |
| H | -2.826339 | 3.014263 | -1.686467 |
| H | -4.406292 | 1.099377 | -1.709254 |
| H | -2.185284 | -0.812627 | -1.866457 |
| H | -4.550064 | -1.505474 | -1.864261 |
| H | -4.760153 | -0.917892 | -0.216205 |
| H | -2.917382 | -3.222485 | -1.045715 |
| H | -4.456206 | -3.383009 | -0.198295 |
| H | -2.548814 | -3.479918 | 1.417447 |
| H | -3.553182 | -2.064839 | 1.724498 |
| H | -0.936856 | -2.121789 | 0.157339 |
| H | -2.692575 | 0.245006 | 0.925703 |
| H | 2.542488 | 1.547508 | 1.979463 |
| H | 2.294029 | -2.185807 | -0.337007 |
| H | 4.495820 | -3.142905 | -0.922772 |
| H | 6.520727 | -1.697735 | -0.963311 |
| H | 6.312845 | 0.723512 | -0.420067 |
| H | 4.106073 | 1.681256 | 0.140296 |
| H | 0.038271 | -0.937189 | 2.120005 |
| H | -1.556961 | -0.857394 | 2.888705 |
| H | -0.744845 | -2.417215 | 2.664621 |
| N | 0.498223 | 1.872775 | 1.820691 |
| H | 0.503364 | 2.477696 | 2.632782 |
| C | -0.222545 | 3.035015 | -1.138548 |
| H | -0.309722 | 3.580713 | -2.084283 |
| H | -0.587534 | 3.691664 | -0.341329 |
| H | 0.839785 | 2.832384 | -0.966400 |

**9**

E -1058.658800

G -1058.277655

0 1

| | | | |
|---|---|---|---|
| C | 1.204565 | -0.923312 | -0.075607 |
| C | 1.952150 | -1.685918 | -1.224447 |
| C | 2.514148 | -0.764063 | -2.276259 |
| C | 2.754175 | 0.534443 | -2.103209 |
| C | 2.440425 | 1.288442 | -0.841240 |
| C | 2.075413 | 0.341770 | 0.334397 |
| C | 1.433182 | 2.430531 | -1.117280 |
| C | 1.077903 | 3.197351 | 0.160569 |
| C | 1.557847 | 1.118896 | 1.582630 |
| C | 0.551746 | 2.236596 | 1.232854 |
| C | 2.748808 | 1.664714 | 2.392624 |
| C | 1.260994 | -1.837297 | 1.161023 |
| C | -0.220384 | -0.531390 | -0.521840 |
| C | -1.129427 | -1.456323 | 1.540741 |
| O | 2.289591 | -2.364817 | 1.553156 |
| O | -0.402080 | -0.067850 | -1.640414 |
| N | 0.087010 | -2.012178 | 1.874092 |
| C | -1.348859 | -0.735069 | 0.413210 |
| C | -2.702968 | -0.225660 | 0.090285 |
| C | -2.883146 | 1.034446 | -0.506770 |
| C | -3.845313 | -0.983451 | 0.399422 |
| C | -4.162681 | 1.526247 | -0.755880 |
| C | -5.125444 | -0.488465 | 0.153659 |
| C | -5.289812 | 0.771636 | -0.423035 |
| C | 1.095895 | -2.804878 | -1.842646 |
| H | 3.371139 | 1.783858 | -0.518118 |
| H | 3.016047 | -0.135776 | 0.632760 |
| H | 1.875620 | 3.109214 | -1.858704 |
| H | 0.529378 | 2.017145 | -1.571143 |
| H | 0.324223 | 3.965060 | -0.055787 |
| H | 1.963112 | 3.728985 | 0.536236 |
| H | 1.036204 | 0.413907 | 2.241925 |
| H | -0.385114 | 1.788937 | 0.881083 |
| H | 0.299694 | 2.785094 | 2.149987 |
| H | 3.386153 | 0.844844 | 2.742790 |
| H | 3.374720 | 2.341072 | 1.800645 |
| H | 2.756685 | -1.234552 | -3.228607 |
| H | 3.172636 | 1.120984 | -2.920445 |
| H | -1.921585 | -1.646552 | 2.257322 |
| H | -2.017542 | 1.624415 | -0.782404 |
| H | -3.729736 | -1.982693 | 0.811752 |
| H | -4.279027 | 2.504766 | -1.214222 |
| H | -5.993105 | -1.095294 | 0.399036 |
| H | -6.286043 | 1.157085 | -0.622337 |
| H | 0.177432 | -2.567483 | 2.716060 |
| H | 0.252196 | -2.397173 | -2.405776 |
| H | 0.707914 | -3.479026 | -1.068892 |
| H | 1.703840 | -3.405878 | -2.528103 |
| H | 2.801178 | -2.171335 | -0.725082 |
| H | 2.399526 | 2.221500 | 3.270241 |

**10**

E -1058.667281

G -1058.28534

0 1

| | | | |
|---|---|---|---|
| C | -3.245400 | 1.329924 | 1.044727 |
| C | -4.055529 | 1.310089 | -0.258383 |
| C | -4.221237 | -0.122689 | -0.779780 |
| C | -2.872871 | -0.841137 | -0.974864 |
| C | -2.070811 | -0.831998 | 0.360896 |
| C | -1.896646 | 0.600958 | 0.916843 |
| C | -0.706414 | -1.467660 | 0.269639 |

| | | | |
|---|---|---|---|
| C | 0.420200 | -0.693520 | 0.094510 |
| C | -0.882898 | 1.368372 | 0.049667 |
| C | -3.081626 | -2.252954 | -1.534774 |
| O | 0.384037 | 0.653215 | -0.028780 |
| C | -0.544767 | 2.726702 | 0.582936 |
| C | -0.768940 | 3.865496 | -0.075465 |
| C | -0.473851 | 5.241201 | 0.446512 |
| C | -0.590783 | -2.891438 | 0.508119 |
| C | 1.751703 | -1.261166 | 0.062084 |
| C | 1.834091 | -2.614453 | 0.238737 |
| N | 0.727286 | -3.380225 | 0.449810 |
| O | -1.515961 | -3.668630 | 0.752802 |
| C | 2.981025 | -0.452622 | -0.124513 |
| C | 4.112687 | -0.701131 | 0.670325 |
| C | 3.067415 | 0.545878 | -1.110245 |
| C | 5.297822 | 0.008679 | 0.476409 |
| C | 4.248737 | 1.259968 | -1.297888 |
| C | 5.370858 | 0.993544 | -0.509180 |
| H | -4.773957 | -0.120231 | -1.728374 |
| H | -3.554389 | 1.922899 | -1.020947 |
| H | -5.036712 | 1.771757 | -0.090967 |
| H | -3.824647 | 0.820835 | 1.827142 |
| H | -3.086927 | 2.359559 | 1.385068 |
| H | -2.294960 | -0.272499 | -1.719975 |
| H | -2.653671 | -1.418334 | 1.083563 |
| H | -1.449668 | 0.521655 | 1.918189 |
| H | -4.827918 | -0.701880 | -0.066955 |
| H | -1.253405 | 1.457610 | -0.977601 |
| H | -2.130784 | -2.730660 | -1.787156 |
| H | -3.693287 | -2.212553 | -2.444614 |
| H | -3.579801 | -2.896948 | -0.803875 |
| H | -0.103277 | 2.740305 | 1.579916 |
| H | -1.203009 | 3.815103 | -1.076069 |
| H | -0.034805 | 5.207879 | 1.449174 |
| H | -1.386690 | 5.850215 | 0.493175 |
| H | 0.224209 | 5.770956 | -0.215327 |
| H | 2.782297 | -3.139094 | 0.205492 |
| H | 0.810668 | -4.379052 | 0.593795 |
| H | 4.052852 | -1.442379 | 1.463179 |
| H | 2.201947 | 0.759264 | -1.726890 |
| H | 6.158825 | -0.199453 | 1.106368 |
| H | 4.294633 | 2.026050 | -2.067669 |
| H | 6.290270 | 1.553619 | -0.657363 |

#### Auto Dock Vina poses and random seeds

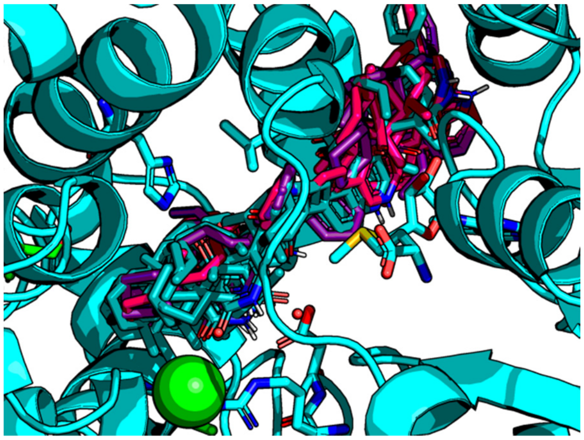

Overlay of all docked ensembles showing entry and exit path to and from active site.

**TS-1**

*random seed:* -1014947296

pose affinity

1 -10.1

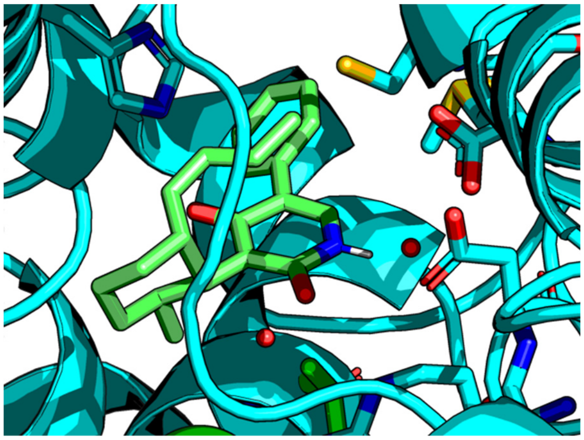

**TS-2**

*random seed:* 1078355739

pose affinity

1 -8.5

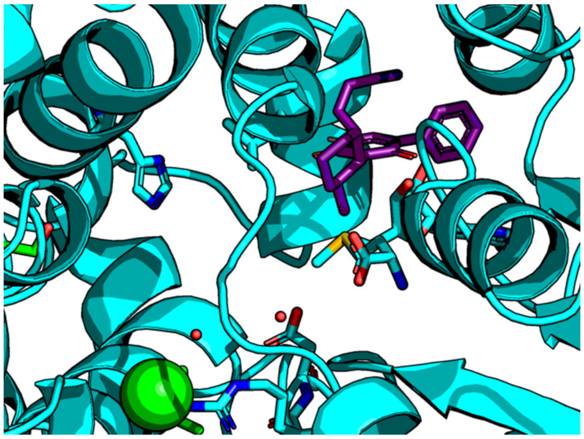

2 -7.8

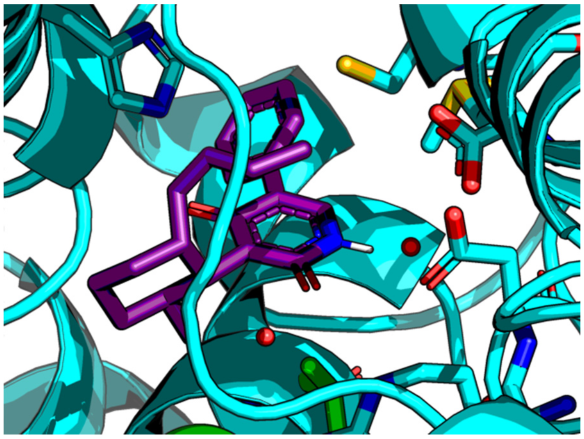

**TS-3**

*random seed:* 2072225248

pose affinity

1 -9.7

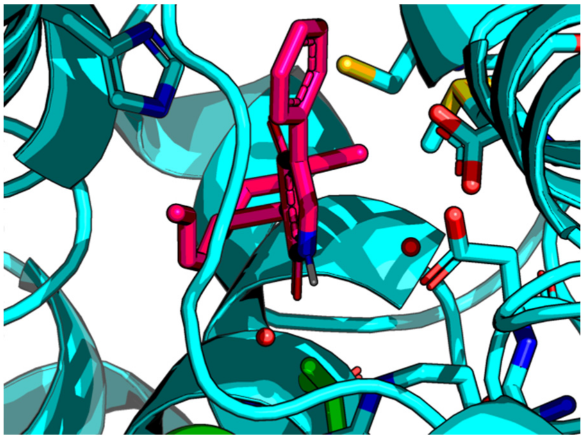

2 -9.4

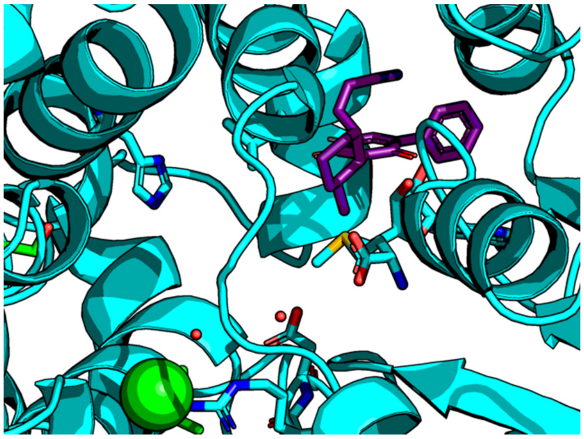

3 -9.2

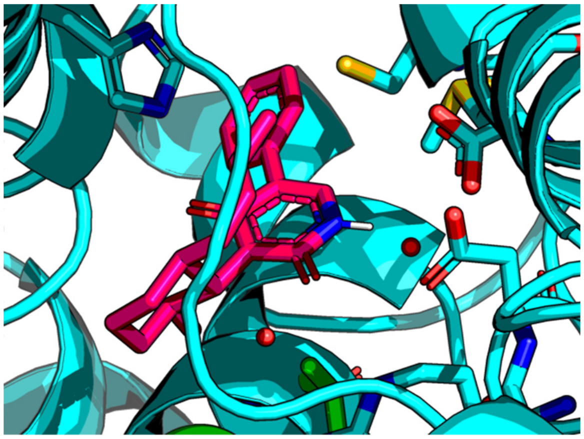

**TS-4**

*random seed:* -1367774985

pose affinity

1 -8.4

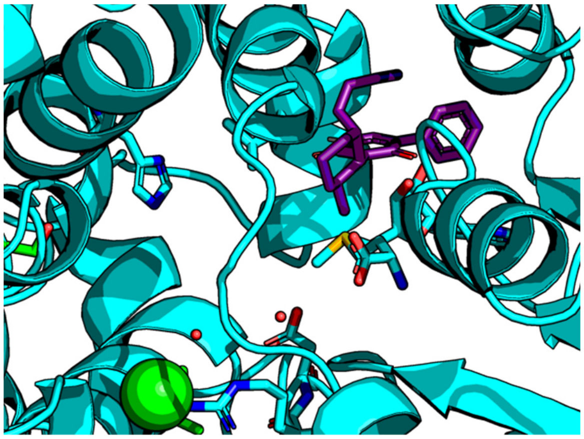

2 -7.5

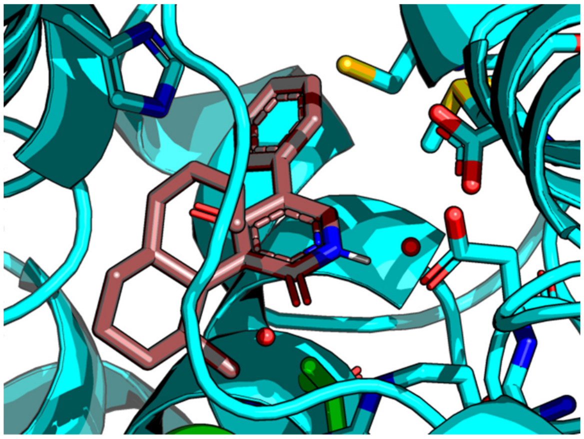

**9**

*random seed:* 309599536

pose affinity

1 -7.0

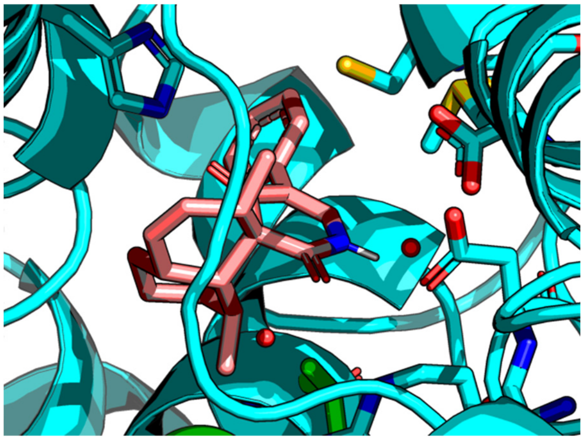
